# Selection, linkage, and population structure interact to shape genetic variation among threespine stickleback genomes

**DOI:** 10.1101/471870

**Authors:** Thomas C. Nelson, Johnathan G. Crandall, Catherine M. Ituarte, Julian M Catchen, William A. Cresko

## Abstract

The outcome of selection on genetic variation depends on the geographic organization of individuals and populations as well as the syntenic organization of loci within the genome. Spatially variable selection between marine and freshwater habitats has had a significant and heterogeneous impact on patterns of genetic variation across the genome of threespine stickleback fish. When marine stickleback invade freshwater habitats, more than a quarter of the genome can respond to divergent selection, even in as little as 50 years. This process largely uses standing genetic variation that can be found ubiquitously at low frequency in marine populations, can be millions of years old, and is likely maintained by significant bidirectional gene flow. Here, we combine population genomic data of marine and freshwater stickleback from Cook Inlet, Alaska, with genetic maps of stickleback fish derived from those same populations to examine how linkage to loci under selection affects genetic variation across the stickleback genome. Divergent selection has had opposing effects on linked genetic variation on chromosomes from marine and freshwater stickleback populations: near loci under selection, marine chromosomes are depauperate of variation while these same regions among freshwater genomes are the most genetically diverse. Forward genetic simulations recapitulate this pattern when different selective environments also differ in population structure. Lastly, dense genetic maps demonstrate that the interaction between selection and population structure may impact large stretches of the stickleback genome. These findings advance our understanding of how the structuring of populations across geography influences the outcomes of selection, and how the recombination landscape broadens the genomic reach of selection.

## Introduction

Biologists have long known that natural populations harbor abundant genetic variation that is distributed heterogeneously across geography (Dobzhansky and Queal 1938; Hubby and Lewontin 1966; Lewontin and Hubby 1966). A recent wave of discoveries reveals that genetic variation is distributed heterogeneously across genomes as well (Turner *et al.* 2005; Begun *et al.* 2007; Hohenlohe *et al.* 2010; Ellegren *et al.* 2012). Are these patterns related? For example, does the manner in which evolutionary processes play out across the geography of organisms influence heterogeneous genomic patterns of genetic diversity? With ever-increasing access to DNA sequence data, evolutionary biologists have come to recognize that selection strongly influences patterns of linked neutral variation (Charlesworth *et al.* 1993; Hahn 2008; Langley *et al.* 2012; Schrider and Kern 2017). Linked positive or purifying selection can affect genetic variation far beyond the causal mutations to the surrounding genomic neighborhood (Elyashiv *et al.* 2016; Schrider and Kern 2017). Indeed, persistent and pervasive linked selection can structure genetic variation during speciation (Burri *et al.* 2015), leading to predictable genome-wide genetic divergence across multiple speciation events (Stankowski *et al.* 2018). Consequently, any process that can influence linkage across the genome, such as variation in recombination, will affect the outcome of linked selection on patterns of genetic variation (Begun and Aquadro 1992; Charlesworth *et al.* 1993; Gillespie 2000).

Interacting with the genomic context of physical linkage is the geographic context over which evolutionary processes play out (Charlesworth *et al.* 1997; Lenormand 2002; Stankowski *et al.* 2017). A single selective sweep in one population can eliminate genetic variation at a selected locus and linked variants (Maynard Smith and Haigh 1974; Fay and Wu 2000; Hermisson and Pennings 2005; Nielsen *et al.* 2005; Cutter and Payseur 2013). However, in nature selective pressures tend to vary across time and space in ways that can maintain variation. (Clausen *et al.* 1941; Endler 1977; Lenormand *et al.* 1999). Temporally fluctuating selection can maintain alleles at intermediate frequencies, as has been observed in ***Drosophila melanogaster*** (Bergland *et al.* 2014) and the yellow monkeyflower, ***Mimulus guttatus*** (Troth *et al.* 2018). Local adaptation in a geographically structured species can lead to the partitioning of variation among populations and the maintenance of alternatively adaptive alleles (Charlesworth *et al.* 2003; Wallbank *et al.* 2016; Nelson and Cresko 2018) as has been observed in insects (Mettler *et al.* 1977; Nosil and Crespi 2006; Van Belleghem *et al.* 2018), plants (Clausen *et al.* 1941; Angert *et al.* 2018; Tavares *et al.* 2018), and vertebrates (Hoekstra *et al.* 2004; Jones *et al.* 2018). Consequently, gene flow can strongly influence the geographic distribution of genetic variation among populations (Wright 1931; Slatkin 1985; Slatkin 1991) and across species boundaries (Pardo-Diaz *et al.* 2012; Fontaine *et al.* 2015). Although genetic introgression from one species to another may impede neutral divergence, it is now recognized as an important source of adaptive variation (Stankowski and Streisfeld 2015; Wallbank *et al.* 2016; Meier *et al.* 2017; Jones *et al.* 2018).

Even in the absence of selection, population subdivision alone may affect the overall abundance of neutral genetic variation within a species. Theoretical models provide useful – but at times contradictory – predictions on this matter, with subdivision either increasing or decreasing expected coalescence times depending on model specifications and assumptions. For example, Slatkin (Slatkin 1991) generalized the island model to demonstrate that population structure increases coalescence times (T_T_) in a structured population by:

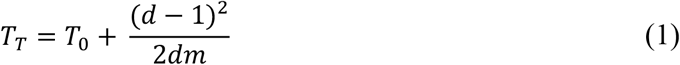

Where d is the number of demes, m is the migration rate, and T_0_ is the expected coalescence time with no population structure (using the nomenclature of Charlesworth *et al.* 2003). Whitlock and Barton (1997), however, used a similar model but allowed deme extinction and variable contributions of each deme to the total population. The result was a decrease in effective population size, N_e_, with increasing subdivision, and thus a decrease in expected levels of genetic variation.

Despite this growing understanding of the general importance of genomic organization of selected loci and geographic context of organisms in the wild on the structuring genetic variation, precise understanding of how they *interact* to produce heterogeneous patterns of genetic variation across the genome is still lacking. Therefore, empirical and simulation studies that examine the joint effects of selection regime, geographic context, and recombination are necessary for fully understanding the effects of these biological factors on genomic patterns of genetic variation in the wild.

Here, we investigate how multiple evolutionary forces interact to shape the genomic and geographic distribution of genetic variation in the threespine stickleback fish, ***Gasterosteus aculeatus***. The threespine stickleback is distributed holarctically in coastal marine, brackish and freshwater habitats. The large marine population has repeatedly given rise to derived freshwater populations, resulting in phenotypic divergence and parallel evolution throughout the species range (Bell and Foster 1994; Cresko *et al.* 2004; Jones *et al.* 2012). When marine stickleback invade freshwater habitats, more than a quarter of the genome can rapidly respond to the action of divergent selection (Hohenlohe *et al.* 2010; Terekhanova *et al.* 2014; Bassham *et al.* 2018). This process largely uses standing genetic variation (Colosimo *et al.* 2005; Schluter and Conte 2009; Roesti *et al.* 2015; Samuk *et al.* 2017) that can be found ubiquitously at low frequency in marine populations (Bassham *et al.* 2018) much of which has likely been maintained for millions of years by significant bidirectional gene flow (Caldera and Bolnick 2008; Kitano *et al.* 2008; Berner *et al.* 2009; Nelson and Cresko 2018).

While previous work has often focused on the genomic targets of selection, we focus here on the processes that maintain and structure variation in regions of the genome physically linked to those targets. We use a combination of population genomics of wild stickleback, forward-time simulations using SLiM (Haller and Messer 2017), and dense genetic maps to support a model whereby differences in population structure between marine and freshwater habitats has led to divergent outcomes of molecular evolution at loci linked to adaptive variants. Our results provide new links between theoretical and empirical evolutionary genetics, new tools for future work in the stickleback system, and new perspectives on the maintenance of genetic variation.

## Methods

### Study populations and natural genetic variation

Wild threespine stickleback were collected from Rabbit Slough (N 61.5595, W 149.2583), Boot Lake (N 61.7167, W 149.1167), and Bear Paw Lake (N 61.6139, W 149.7539). Rabbit Slough (RS) is an offshoot of the Knik Arm of Cook Inlet and is known to be populated by anadromous populations of stickleback that are stereotypically marine in phenotype and genotype (Figure 1; Cresko *et al.* 2004). Boot Lake (BT) and Bear Paw Lake (BP) are both shallow lakes formed during the end-Pleistocene glacial retreat approximately 12 thousand years ago. Fish were collected in the summers of 2009 (RS), 2010 (BP), and 2014 (BT) using wire minnow traps and euthanized ***in situ*** with Tricaine solution. Euthanized fish were immediately fixed in 95% ethanol and shipped to the Cresko Laboratory at the University of Oregon (Eugene, OR, USA).

**Figure 1.**
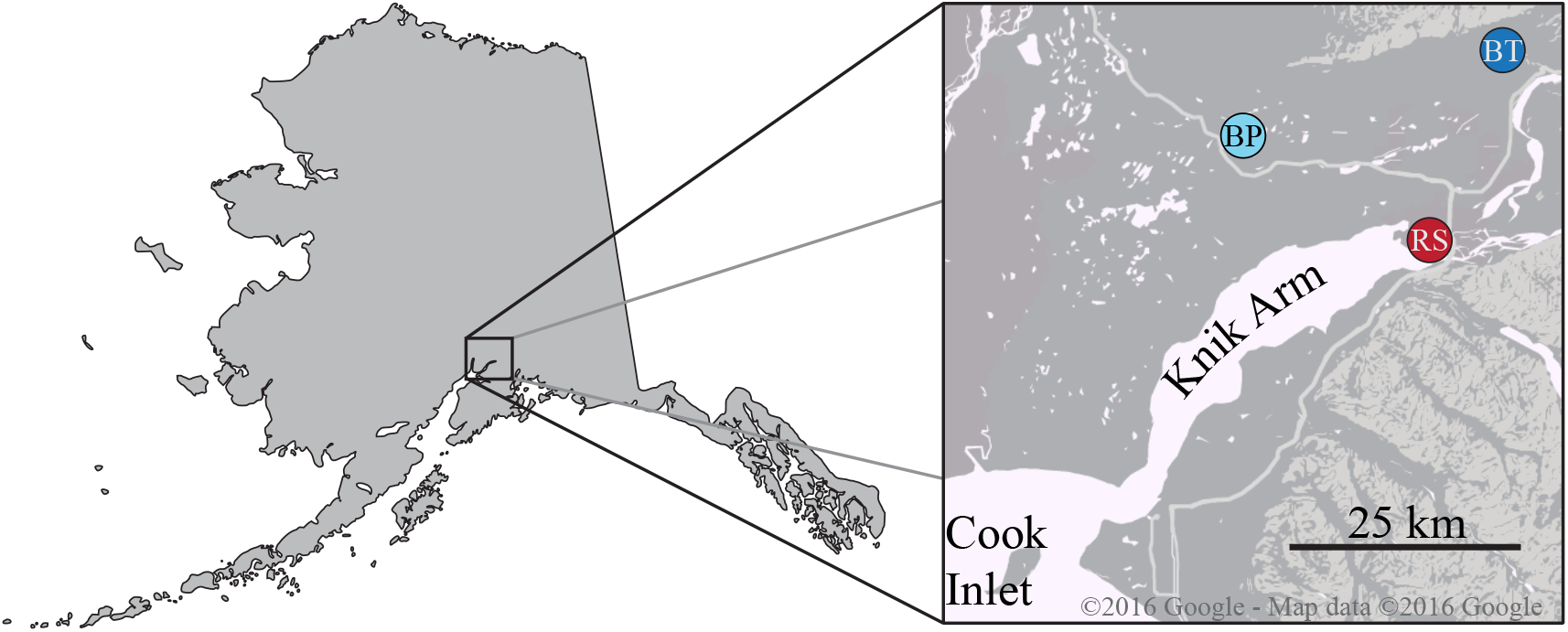
Sampling locations of marine and freshwater stickleback populations from southcentral Alaska. Inset: Geography of Cook Inlet near Anchorage, AK. Marine stickleback were sampled from Rabbit Slough (RS, red). Freshwater stickleback were sampled from Boot Lake (BT, dark blue) and Bear Paw Lake (BP, light blue). All areas in the inset were glaciated during the Late Wisconsin glaciation until ~10 thousand years ago.

We generated restriction site-associated DNA (RAD) libraries of five fish each from RS and BT and four fish from BP. Genomic DNA was isolated from ethanol-preserved fin clips by proteinase K digestion followed by DNA extraction with solid phase reversible immobilization (SPRI) beads. We created RAD libraries using the single digest and shearing method of Baird et al (Baird *et al.* 2008) with the modifications of Nelson and Cresko (Nelson and Cresko 2018). Genomic DNA from each fish was digested with *PstI*-HF (New England Biolabs) and ligated to Illumina P1 adaptors with 6 bp inline barcodes. All barcodes differed by at least two positions, allowing for recovery of sequence reads with single errors in the barcode sequence. Ligated samples were then multiplexed at approximately equimolar concentrations and mechanically sheared via sonication to a fragment range of ~200-800 bp. Sheared DNA was size selected by extraction from a 1.25% agarose gel to generate a narrow insert size range of 425 bp to 475 bp. This size range allowed consistent overlap of paired-end Illumina reads for the construction of local contigs surrounding restriction enzyme cut sites. We then ligated Illumina P2 adaptors to the size-selected libraries and amplified P1/P2-adapted fragments with 12 cycles of PCR using Phusion-HF polymerase (New England Biolabs). RAD libraries were then sequenced in a single lane on an Illumina HiSeq 2500 to generate paired-end 250-bp sequence reads. All libraries generated for this study were sequenced at the University of Oregon’s Genomics and Cell Characterization Core Facility (GC3F: http://gc3f.uoregon.edu).

We used the Stacks analysis pipeline to process RAD sequence read pairs and call SNPs (Catchen *et al.* 2011; Catchen *et al.* 2013b). Raw reads were first demultiplexed without quality filtering using process_radtags, and then quality filtered using process_shortreads. This allowed for read trimming, rather than strict removal, if quality decreased toward the end of the first-end read. Overlapping read pairs were then merged using fastq-join (Aronesty 2011), allowing for up to 25% of bases in the overlapping region to mismatch, and the resulting contigs were trimmed to 350 bp. Any read pairs that failed to merge, or were shorter than 350 bp, were removed from further analysis. This step was required for processing reads through the Stacks pipeline. Below, a “locus” refers to the combined sequence assembled from a single restriction site in the stickleback genome.

All polymorphisms were called relative to the threespine stickleback reference genome v1.0 (Jones *et al.* 2012), using the updated scaffolding of Glazer, et al. (2015). Trimmed contigs were aligned to the reference using bbmap with the most sensitive settings (‘vslow=t’; http://jgi.doe.gov/data-and-tools/bbtools/). We then used the Stacks core pipeline to identify read stacks, call SNPs, and identify alleles and haplotypes based on genomic alignment (pstacks and cstacks); find homologous RAD tags across individuals (sstacks); and catalog biologically plausible haplotypes based on within- and among-individual haplotype variation (populations). We required that a RAD tag be present in all three populations and in at least four fish in each population.

We used the program PHASE (Stephens *et al.* 2001; Scheet and Stephens 2006) to combine sequence information from both RAD tags at a ***PstI*** cut site and generate phased haplotypes at each RAD locus. We wrote custom Python scripts to identify all unique haplotypes at each pair of RAD tags and code them as alleles at a single, multiallelic locus. We required that each individual included in this analysis was genotyped at both RAD tags. Loci containing individuals only genotyped at a single RAD tag were removed from further analysis. RAD haplotypes at each locus represent 696 bp of contiguous genomic sequence, giving us high-quality estimates of sequence diversity and divergence even with our relatively small populationlevel sample sizes (Nei 1987, chapter 13; Cruickshank and Hahn 2014; Nelson and Cresko 2018). Sequencing of wild stickleback resulted in 57,992 RAD loci distributed across all 21 threespine stickleback chromosomes, averaging 7,514 bp between adjacent RAD loci.

### Population genetic statistics

The scripting language R version 3.5 (R Core Team 2016) was used for all downstream data analysis. We estimated differentiation among threespine stickleback populations (all pairwise combinations) and among ecotypes (combined freshwater ponds versus RS) using a haplotype-based F_ST_ (equation 3 in Hudson *et al.* 1992) implemented in the R package ‘PopGenome’ (Pfeifer *et al.* 2014). We calculated π per site within and among populations and absolute sequence divergence (d_XY_) at each RAD locus by calculating pairwise distances between all RAD haplotypes with the R package ‘ape’ (Paradis *et al.* 2004; Popescu *et al.* 2012).

Previously, we detected patterns of reciprocal monophyly between marine and freshwater haplotypes using maximum clade credibility trees generated in BEAST v1.7 (Drummond and Rambaut 2007; Drummond *et al.* 2012; Nelson and Cresko 2018; Suchard *et al.* 2018). Tree topologies for all RAD loci were inferred from MCMC runs of 10^6^ states with 10% burnin periods. We used blanket priors and parameters across all RAD loci, including a coalescent tree prior and the GTR+Γ substitution model. Monophyly of haplotypes from each population (RS, BT, BP) and each habitat (marine, freshwater) was assessed using the is.monophyletic() function of the R packages ‘ape’. Here, we use topological classifications that we inferred previously and designate gene trees with reciprocally monophyletic marine and freshwater haplogroups (1,129 of 57,992 RAD loci) as ‘divergent’ loci (Figure 2B, see also Nelson and Cresko 2018).

**Figure 2.**
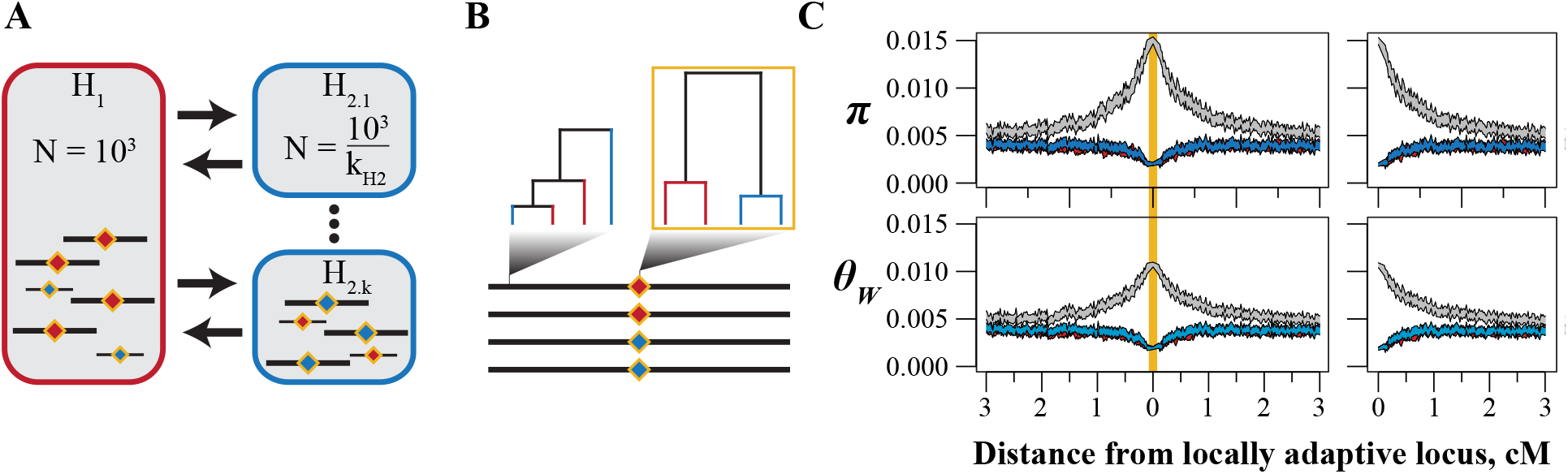
The structure of simulations to test the effects of population structure on genetic variation in a model of local adaptation. **A:** Habitat H_1_ (red) (: red, H_2_: blue) consists of a single deme while habitat H_2_ (blue) consists of *k* demes. We adjust the size of each deme in H_2_ so that the total census size of each habitat is constant across simulations. Diamonds represent a single locus with locally adaptive alleles. **B:** We assessed genetic diversity at the locally adaptive locus and in non-overlapping windows. We used the pattern of reciprocal monophyly (outlined in gold) to classify RAD loci as ‘divergent’ in the threespine stickleback population genomic dataset. (See Methods and Nelson and Cresko [2018]). **C:** Genetic diversity within and among allelic classes in a simulation where *k*_H2_ = 1. Shown are 95% confidence intervals of nucleotide diversity (π) and Watterson’s θ after 10N generations of selection. Because patterns of variation are symmetrical about the locus under selection, we present ‘folded’ curves (panel C, right) throughout this work.

### Forward simulations using SLiM

We used forward simulations implemented in SLiM (Haller and Messer 2017) to model the effects of selection, linkage, and population structure in a manner that reflects the stickleback metapopulation structure (Figure 2). We simulated a metapopulation of 2000 diploid individuals and a genome consisting of a single chromosome with a genetic length of ten centiMorgans (cM). The chromosome contained 50 kb of freely recombining sequence (recombination rate 1×10^-6^ per bp) on either side of a 2 kb nonrecombining ‘core’ containing the locally adaptive locus. Per-base mutation rate was kept constant at 5×10^-7^, resulting in a population-scaled mutation rate (4Nμ = 0.004) aligned with genome-wide estimates of genetic diversity in stickleback (Hohenlohe *et al.* 2010; Nelson and Cresko 2018). The general form of the simulations was as follows:

1. Initialize population H_1_
  a. N_H1_ = 1000 diploids
  b. evolve for 10,000 generations
2. Create *k* new populations, collectively “H_2_”
  a. each of size 1000/*k*
  b. set migration rate m to H_1_
  c. evolve for 10,000 generations
3. Introduce mutation as single copy into H_2_
  a. Deleterious in H_1_, advantageous in H_2_
  b. Run selection for [4,10,20]N generations
  c. (If locally adaptive mutation is lost, go to [3])
4. End simulation

Simulations included three phases: (1) a burn-in without population structure, (2) a burn-in after creating population structure, and (3) a selection phase contingent on establishment of a locally adaptive allele. The first burn-in began with a single panmictic population of size N_H1_ = 1000 and no sequence variation. We evolved this population for 10N_H1_ (10,000) generations. The second burn-in of 10N_H1_ generations began by creating *k*_H2_ new populations, each of size N_H1_/k_H2_, by sampling variation from the existing population (Figure 2), where *k*_H2_ ranged from one to twenty-five. We set bidirectional migration rates equivalent to one migrant per generation between the existing population, now designated habitat H_1_, and each new population, now collectively referred to as habitat H_2_. We chose this migration structure to reflect the stickleback metapopulation, where freshwater stickleback populations are thought to be derived most often from marine ancestors, and because gene flow among freshwater populations primarily occurs through the marine population, especially across broader geographic scales.

To start the selection phase, we introduced a mutation at the center of the ‘core’ of a single chromosome in habitat H_2_ that is advantageous in H_2_ and deleterious in H_1_. Progression through the selection phase was conditional on establishment of the locally adaptive mutation: if the mutation was lost from the metapopulation, the simulation restarted at the end of the second burn-in.

Genetic diversity among chromosomes carrying H_1_- and H_2_-adaptive alleles was assessed at the end of the selection phase. We sampled 10 chromosomes of each allelic state and calculated nucleotide diversity (π) and Watterson’s θ (θ_W_) within and among chromosomes of the two allelic states in non-overlapping 250 bp (0.025 cM) bins. We include results from θ_W_ because this statistic is based on the number of segregating sites and is therefore more reflective of the abundance of sequence variation and less affected by shifts in allele frequencies. We fold the calculations of genetic diversity about the locally adaptive locus to emphasize the effects of evolutionary forces on π at a given distance from the locus (Figure 2C). These summaries mirror our estimates of genetic variation on stickleback chromosomes and results from theoretical work on the effects of local adaptation on linked variation (Charlesworth *et al.* 1997).

### Laboratory crosses and genetic mapping

To compare how heterogenous genomic divergence on the physical map is distributed across the genetic map, we generated mapping families from laboratory lines of fish derived from the Boot Lake (BT) and an F_1_ hybrid female (hereafter F_1_) produced from a cross between a BT female and a Rabbit Slough (RS) male. These crosses allowed us to examine variation in the recombinational landscape within and among chromosomes in distinct, evolutionarily relevant genetic backgrounds. We also generated a genetic map of a RS-derived male (Figure S1) but, because ecotype was confounded with sex (Sardell *et al.* 2018), we limit our discussion to the BT and F_1_ maps.

All maps were constructed using a pseudo-testcross design, which takes advantage of existing heterozygosity in outbred individuals without the need to generate inbred lines or F_1_ mapping parents. To generate the BT mapping family, we manually crossed unrelated lab-reared individuals. We mapped the F_1_ female by backcrossing it to a BT male. All progeny were raised to 14 days post-fertilization, euthanized with MS-222 (Sigma Aldrich), and fixed in 95% ethanol. We extracted genomic DNA from whole progeny and from pectoral and caudal fins from all parents using proteinase K digestion (Qiagen) followed by DNA purification with SPRI beads.

RAD genotyping of progeny and parents was used to identify segregating haplotypes using a RAD-seq protocol similar to that described previously but using the restriction enzyme *SbfI*. RAD-seq data from all mapping crosses were processed with the Stacks analysis pipeline (Catchen *et al.* 2013b). We demultiplexed and quality filtered sequences with process_shortreads and aligned them to the stickleback reference genome with GSNAP (Wu and Nacu 2010). We used ref_map.pl to identify RAD tags and call genotypes. The Stacks component program genotypes was used to identify segregating markers for export to genetic mapping software. We specified a minimum coverage of 3x to call individual genotypes and required that a marker be genotyped in at least 50% of progeny. Below, we use the term ‘RAD marker’ to refer to a RAD tag with segregating haplotypes.

Below, we present genetic maps for the female parent from each cross (Table 1; Table S1). By conducting pseudo-testcrosses, we identified polymorphic RAD markers segregating in all mapping parents. However, to investigate the genome-wide recombination landscape, as well as relationships between recombination rate and natural levels of polymorphism and divergence, we restricted our analysis to parents of the same sex for which we observed segregating markers on all 21 chromosomes with no gaps of more than 1 mega-base pairs (Mbp) between adjacent markers.

**Table 1.** Genetic map statistics for the three maps used in this study. *map length given in centiMorgans (cM). †Genomic coverage is a percentage of the reference genome assembled onto chromosomes (436.6 Mb).

| Family | progeny | markers | map length* | genomic coverage† | cM/Mb |
| --- | --- | --- | --- | --- | --- |
| Boot Lake (BT) | 189 | 6041 | 2255 | 0.993 | 5.20 |
| BT subset | 94 | 6169 | 2396 | 0.993 | 5.52 |
| F<sub>1</sub> hybrid | 102 | 6669 | 1878 | 0.995 | 4.33 |
| F<sub>1</sub> subset | 94 | 7678 | 1958 | 0.995 | 4.51 |

We estimated genome-wide recombination rates between RAD markers under the assumption of collinearity between all genetic maps and the stickleback reference genome (Glazer *et al.* 2015) (with some exceptions, see below). We used the mapping software Lep-MAP2 (Rastas *et al.* 2015) to estimate map positions of RAD markers with the marker order fixed to the aligned positions on the reference genome. Known marker orders increased throughput of mapping iterations and reduced the impact of genotyping errors on recombination rate estimation.

While fixed marker orders do not explicitly allow the detection of structural variation among genomes — such as chromosomal inversions that are known to exist among stickleback populations — discrepancies in the estimated map do provide indirect, and correctable, evidence of changes to map order (Figure S2). For example, inversions in genetic map order relative to the reference order spuriously inflate recombination rates between markers closely flanking inversion breakpoints when inverted segments are forced into the wrong orientation. This is because genetic map distances are estimated independently while the physical distance between markers is drastically underestimated. The observed jumps in map distance on either side of the inversion are equal to each other and to the total map length of the inversion. Reversing the marker order within the inversion removes these artifacts and reduces the overall map length of the region. We observed and corrected this artifact for a single previously known inversion on chromosome 21 (Figure S1, Figure S2, Jones *et al.* 2012). Two other known inversions on chromosomes 1 and 11 were too small to create artifacts in our data.

### Recombination-polymorphism correlations

We employed three methods to investigate the relationships between the recombinational landscape and patterns of polymorphism within and among natural populations. To compare our results to other studies in stickleback (Samuk *et al.* 2017), we quantified genome-wide correlations among recombination rate and population genetic statistics. We divided the stickleback genome into non-overlapping windows and calculated average recombination rates (in centiMorgans per mega-base pair, cM/Mbp), sequence diversity (π_BT_, π_RS_, and π), and genetic divergence (F_ST_, d_XY_) in each window. We calculated F_ST_ between RS and BT as a measure of recent genetic differentiation and d_XY_ between RS and the combined freshwater populations as a measure of long-term genetic divergence (Nelson and Cresko 2018). RAD marker density made estimates of local recombination rates less reliable and more variable when using small window sizes (e.g. 100-kbp windows; Figures S3 and S4). As a result, we show results from 1-Mbp genomic windows. We used nonparametric correlations to test for correlations between variables because the distributions of most variables lacked normality even using standard data transformations. Below we present Spearman’s rank order correlations. Kendall’s tau and parametric linear models gave qualitatively similar results (Table S2).

Genomic heterogeneity exists not only in the proportion of the genome affected by divergent selection but also in how genetic variation and divergence are clustered within the genome. Marine-freshwater genomic divergence in stickleback is clustered into few, large regions that can encompass most of the length of a chromosome. We sought to directly compare the genomic distributions of population genetic statistics along the physical genome and on the genetic maps we constructed. We used a windowing approach that allowed direct comparisons across maps despite differing numbers and distributions of markers among genetic maps. Using the R package ‘ksmooth’, we binned each chromosome into equally sized intervals, the number of which we set equal to the number of segregating RAD markers on the genetic map with fewest markers. For each interval, we calculated genetic position from each laboratory cross and F_ST_ between RS and BT populations (Hohenlohe *et al.* 2010; Nelson and Cresko 2018) within a 250-kbp normally distributed kernel. We also imputed the genetic positions of all divergent loci using the lm() function in R: we found the nearest flanking RAD markers in each mapping cross and predicted the genetic position of the divergent locus assuming a constant recombination rate between the flanking markers. With these approaches we were able to make direct comparisons between recombination within and among genetic maps from different genetic backgrounds and patterns of polymorphism in the populations from which the laboratory lines were derived.

### Data availability

Raw sequence data will be made available on NCBI under BioProjects PRJNA429207 and PRJNAXXXXXX. RAD sequences for mapping families are available on the Sequence Read Archive, BioSamples SAMN10498162-10498548. Scripts and processed data will be available on FigShare and are immediately available on GitHub at https://github.com/thomnelson/linkedvariation.

### Animal care and compliance

Treatment of animals followed protocols approved by the University of Oregon Institutional Animal Care and Use Committee (IACUC).

## Results

### Chromosomes from freshwater but not marine stickleback contain abundant linked variation

To examine patterns of variation linked to loci under selection, we first partitioned RAD loci from the population genomics dataset into those with evidence of complete marine-freshwater lineage sorting (‘divergent’, with allelic states ‘marine’ and ‘freshwater’, see Figure 2B), those on the same chromosome as a divergent locus (‘linked’), and those on chromosomes without divergent loci (‘unlinked’). At divergent loci, average sequence diversity was partitioned almost entirely among chromosomes carrying marine and freshwater alleles; overall π averaged 0.0067 per site (Figure 3A, black-filled diamond, genome-wide mean = 0.0042); average d_XY_ between marine and freshwater alleles was 0.0124, nearly three-fold higher than the genome-wide average (0.0044). In contrast, and as expected at locally adaptive loci (Figure 2C), π was substantially reduced within allelic states; π among marine alleles averaged 0.0012 per site (Figure3B, red-filled diamond) while average π among freshwater alleles at divergent loci was 0.0015 per site (Figure3C, blue-filled diamond).

**Figure 3.**
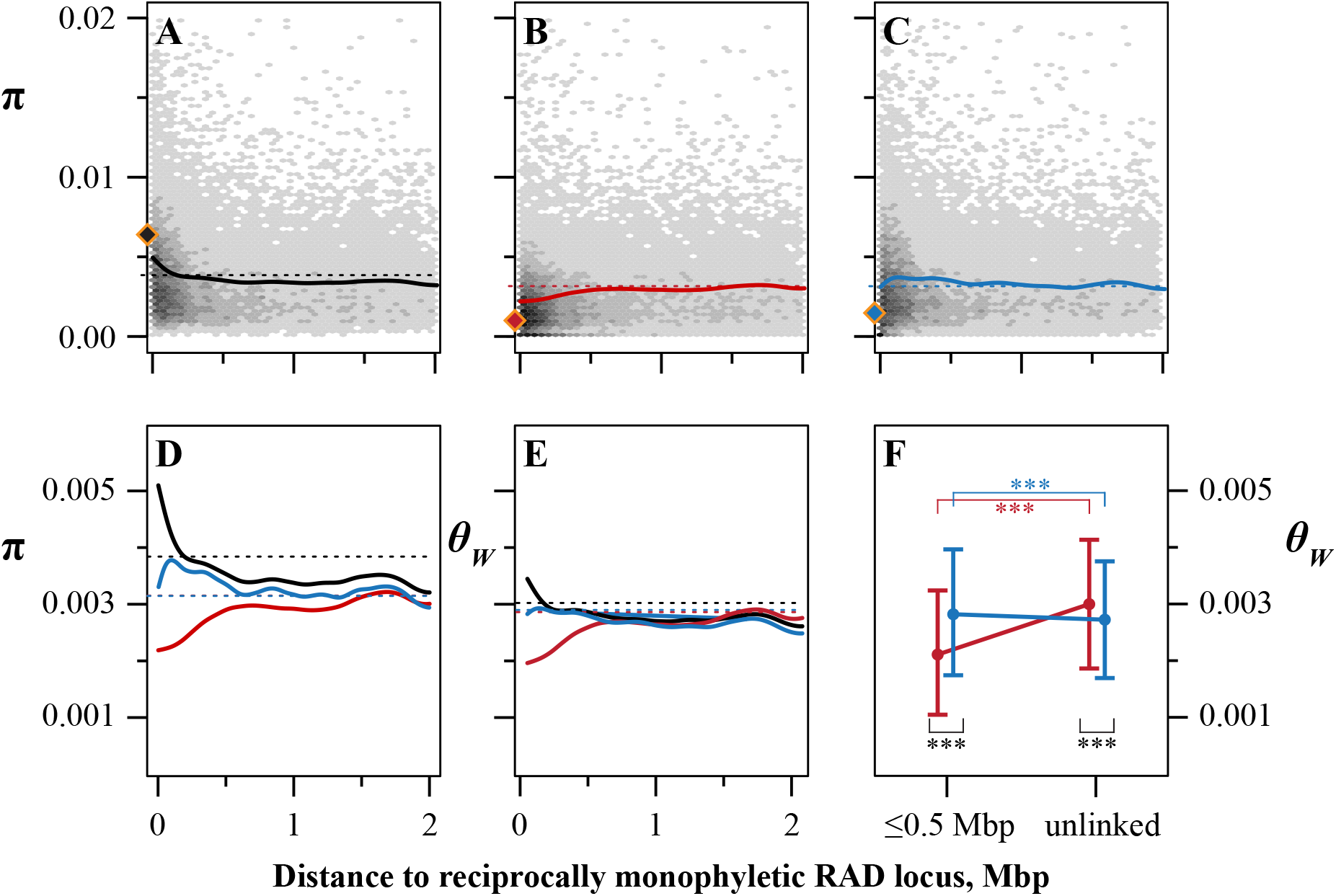
Patterns of sequence variation differ on marine and freshwater stickleback chromosomes. **A-C:** Genome-wide sequence diversity (π) as a function of the distance from a RAD locus with reciprocally monophyly of marine and freshwater haplotypes. Diamonds show average sequence diversity among reciprocally monophyletic RAD loci. Density of RAD loci at x distance from the nearest reciprocally monophyletic locus with y level of diversity is shown in shades of gray. Bold lines are smoothed splines, dashed lines are genome-wide means. **A:** Sequence diversity in the combined dataset (marine [RS] + freshwater [BT and BP]). **B:** Sequence diversity on chromosomes carrying marine haplotypes. **C:** Sequence diversity on chromosomes carrying freshwater haplotypes. **D:** Smoothed splines as in A-C. **E:** Sequence diversity of RAD loci within 1 Mb of a reciprocally monophyletic RAD locus and on chromosomes not carrying a reciprocally monophyletic locus (‘unlinked’). Data are means ± 1 SD. *** p < 0.001. **F:** Splines as in **A-D** of Watterson’s θ.

Patterns of linked variation among habitats and on marine chromosomes followed the expectations from simulated local adaptation (Figure 2C, Figure 3A,B). Sequence diversity at linked loci was highly correlated with proximity to a divergent locus (Figure 3A-C, Spearman’s ρ: all p-values ≤ 10^-10^). When all populations were combined, π at linked loci decreased sharply in the first approximately 250 kb from a divergent locus (Figure 3A,D), and linked loci in closest proximity to divergent loci were nearly as polymorphic as those directly impacted by divergent selection. As expected under local adaptation, π among chromosomes sampled from marine RS stickleback was lowest in close proximity to divergent loci (Figure 3B), and a substantial fraction (12%) of RAD loci within 250 kb of a divergent locus showed no variation at all on these chromosomes (π_RS_ = 0; 1651/13762 loci).

In stark contrast, diversity among freshwater chromosomes showed a proximity effect that was distinct from either RS or the combined populations (Figure 3C-E). Rather than being lowest near divergent loci, diversity actually increased with proximity to a divergent locus, peaking approximately 200 kbp away on average before reversing direction. This pattern persisted using either π or Watterson’s theta (θ_W_), indicating that the density of segregating sites on freshwater chromosomes is highest near divergent loci and that the signal we observe is not simply due to a greater abundance of intermediate frequency alleles. We also note that this increase in diversity qualitatively persisted within both freshwater populations individually (Figure S5).

Linkage to divergent loci, therefore, was associated with opposing effects on genetic variation in stickleback: decreasing it among marine chromosomes while increasing variation among freshwater chromosomes (Figure 3E). Chromosomes with no evidence of divergent selection had a slightly higher density of segregating sites in RS than the combined freshwater populations (Figure 3E, Table 2), though π was indistinguishable (Figure S6). However, within 500 kbp of a divergent locus, genetic diversity (π and θ_W_) was greater among freshwater populations than in RS (Figure 3E, Table 2, Figure S6; population*proximity interaction p ≤ 10^-10^).

**Table 2.** ANOVA results for proximity-by-ecotype analyses. F statistics are from ANoVAs on untransformed values, ***p ≤ 10^-10^. For Tukey’s Honest Significant Difference (HSD), ‘+’ and ‘−’ indicate an increase or decrease in diversity, respectively, compared to variation at unlinked loci on marine (RS) chromosomes. ‘linked’ indicates loci ≤ 500 kb from a divergent locus, as in Figure 3.

| | $F_{1,34068}$ | | | Tukey's HSD vs $RS_{unlinked}$ | | |
| --- | --- | --- | --- | --- | --- | --- |
| | ecotype | linkage | ecotype*linkage | $RS_{linked}$ | $FW_{linked}$ | $FW_{unlinked}$ |
| $\pi$ | 858.8*** | 62.5*** | 382.8*** | - | + | = |
| $\theta_w$ | 343.4*** | 149.6*** | 358.9*** | - | - | - |

### Population structure maintains linked variation on simulated chromosomes

To determine how population structure within the freshwater habitat influences patterns of linked variation in stickleback, we conducted forward simulations of chromosomes under a model of local adaptation with migration using SLiM v2.6 (Haller and Messer 2017, Figure 2). In simulations with two habitat types, each composed of a single, panmictic population (Figure 4, column 1), total genetic diversity was highest at and adjacent to the locally adaptive locus (Figure 2C). Within allelic classes, diversity was lowest in proximity to the locally adaptive locus and recovered equally within both allelic classes with increasing recombinational distance from the selected locus (Figure 2C; Figure 4). This scenario is essentially the same as that presented by Charlesworth, Nordborg, and Charlesworth (1997) and the results are qualitatively the same.

**Figure 4.**
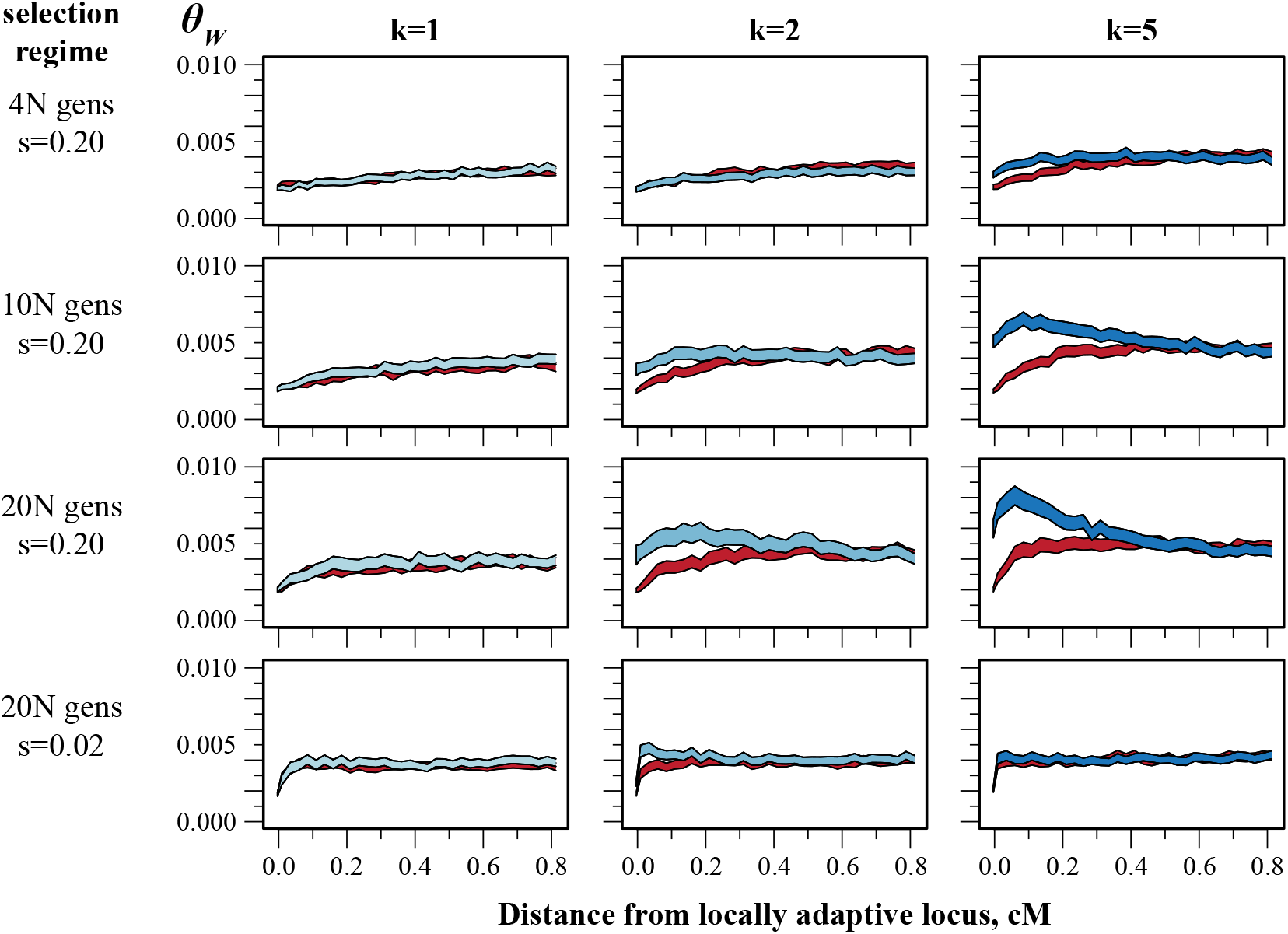
Asymmetric population structure generates asymmetric patterns of linked variation on simulated chromosomes. Simulations were performed as described in Methods and Figure 2. Bands are 95% confidence intervals of Watterson’s θ at a given distance from a locally adaptive locus (red: θ_W_ on chromosomes carrying H_1_-adaptive allele, blue: θ_W_ on chromosomes carrying H_2_-adaptive allele). Columns show the effect of increasing population structure (*k*) in habitat H_2_. Rows show the effect of increasing the length of the selection phase of the simulations. Rows 1-3 show results under strong selection (*s* = 0.20). Row 4 shows results for simulations identical to row 3 but with moderate selection (*s* = 0.02). All simulations were performed using a migration rate, *m*, of one migrant per generation between habitat H_1_ and each population of habitat H_2_. Total diversity (see Figure 2) is excluded to highlight variation within each allelic class.

Adding to these simulations population structure informed by the stickleback system led to previously undocumented results. When the second habitat (H_2_) – representing the freshwater stickleback habitat – consisted of two or more demes (Figure 2A; Figure 4, columns 2 and 3) the additional structure maintained greater variation exclusively on H_2_-adaptive (H_2_^+^) chromosomes, an effect dependent on the duration and strength of selection and the number of demes. Under strong selection (s = 0.20), population structure had a barely noticeable effect on variation until the length of the selection phase was on the order of the neutral coalescence time (~9NH generations, Slatkin 1991; Charlesworth *et al.* 2003). Beyond this point, any amount of population structure resulted in higher levels of variation on H_2_^+^ than H_1_^+^ chromosomes within ~0.2 cM from the selected locus. Moderate selection (s=0.02) was generally ineffective at altering among-habitat levels of variation, though we did observe a modest effect on H_2_^+^ chromosomes immediately adjacent to the selected locus.

Increasing population structure in H_2_ led to substantial levels of variation near the selected locus (Figure 6, column 3; Figure S7). When H_2_ consisted of five demes, genetic variation on H_2_^+^ chromosomes was greatest ~0.1 cM from the selected locus even though levels of variation among allelic states remained indistinguishable in distal regions of the chromosome. Greater subdivision of H_2_ first accentuated the accumulation of variation (5 ≤ *k* ≤ 15, Figure S7) but eventually attenuated it when *k* ≥ 20. While we did fully not explore this attenuation, we note that we held both *N* and *m* (the average number of migrants per generation) constant across simulations; higher values of *k* were therefore accompanied by greater total migration between habitats and stronger within-deme drift in H_2_.

The interaction we observed between selection and geographic structure in simulated populations closely mimicked the patterns found in stickleback, but the extent of this effect seemed limited. Even in the case of strong selection (s=0.20) and substantial population structure, increases in genetic diversity extended less than 0.5 cM away from the locus under selection; maximal levels of variation required even tighter linkage. We therefore examined how recombination rate varies across stickleback genomes to better understand the extent to which selection may influence linked genetic variation.

### The recombination landscape varies among individuals

RAD sequencing of both mapping crosses resulted in over 6000 segregating markers, with markers averaging 70 kbp apart on the BT map and 56 kbp apart on the F_1_ map (Table 1). Mean per-locus sequencing depth was 35x and 23x for the BT and F_1_ families, respectively (range: BT=[28x,57x], F_1_=[10x,90x]; Table S1). RAD markers were consistently genotyped in over 90% of progeny (BT: mean=98%, range=[95%,99%]; F_1_: mean=91%, range=[57%,99%]).

Patterns of recombination were generally consistent between the genetic maps (Figure 5, Figure S1). As has been described previously (Roesti *et al.* 2013; Glazer *et al.* 2015), recombination on the larger metacentric chromosomes in the stickleback genome (e.g chromosomes 4 and 7) was biased toward chromosome ends, with little recombination occurring across central, presumably centromeric, regions. Recombination rates across a number of the smaller chromosomes, in contrast, was typically highest toward one end (Figure 5, chromosome 15; Figure S1).

**Figure 5.**
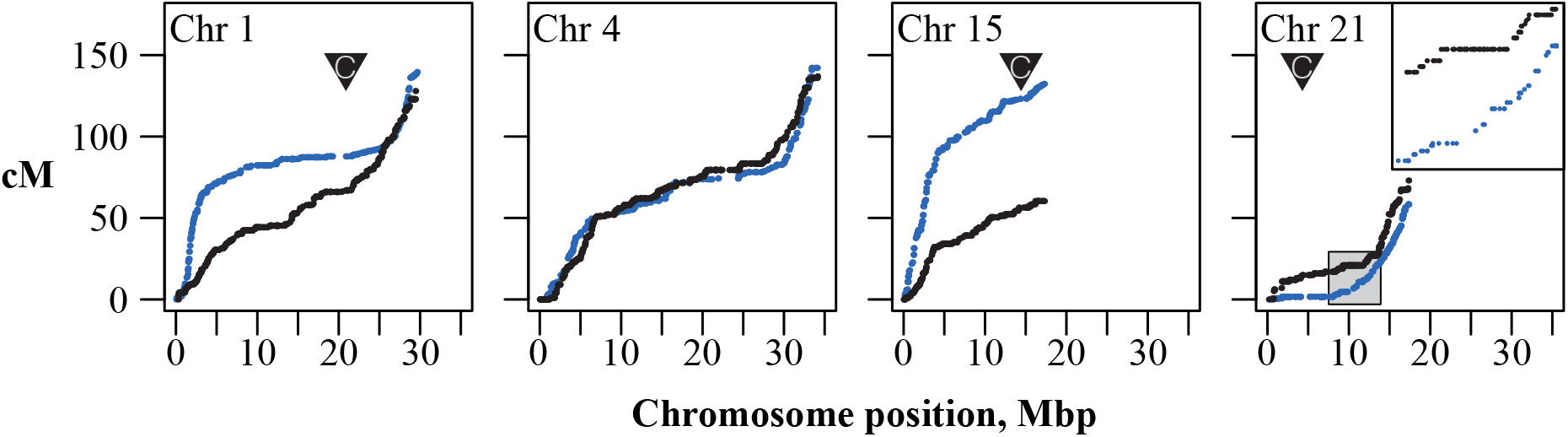
The recombination landscape varies within and among genetic maps of BT and F_1_ hybrid threespine stickleback. Each point represents a RADseq-based marker segregating within a mapping family. Blue: BT, black: F_1_. Inverted triangles represent the locations of centromeric repeats, where known, from Sardell et al (2018). The inset on chromosome 21 shows suppression of recombination due to inversion heterozygosity in the F_1_. Genetic maps for all chromosomes and for the marine (RS) map are given in Figure S1.

**Figure 6.**
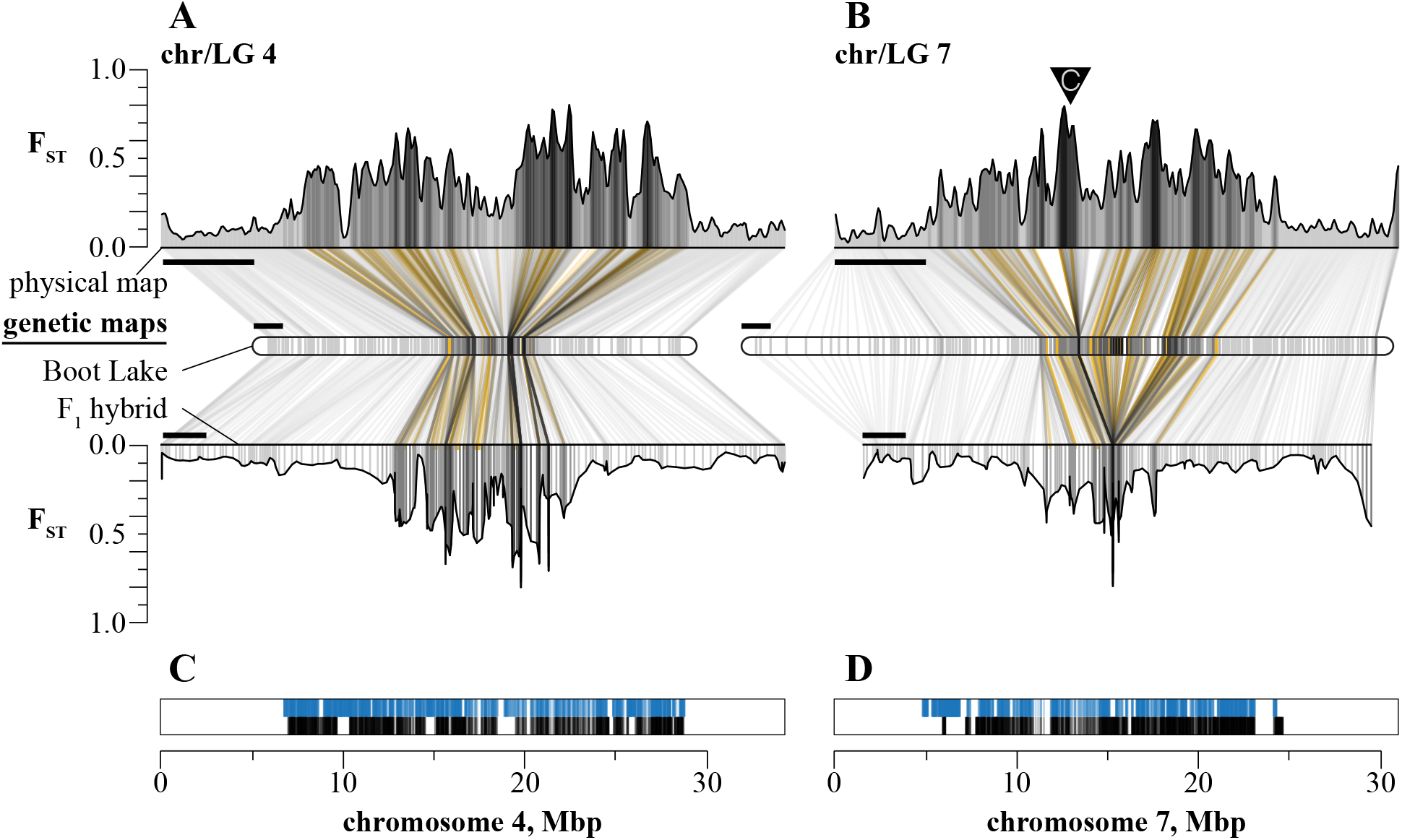
The recombination landscape extends to genomic reach of divergent selection. **A-B:** RS-BT F_ST_ scans were performed on the physical map (top, scale bars = 5 Mbp) and imputed onto the freshwater and F_1_ hybrid genetic maps (middle and bottom, scale bars = 10 cM) for chromosomes 4 and 7. Lines connect evenly-spaced windows on the physical map to the imputed positions on each genetic map and are colored by the kernel-averaged F_ST_ for that window. Positions of divergent RAD loci are shown in gold. Inverted triangle represents the position of centromeric repeats (Sardell et al 2018). **C-D:** physical positions of all RAD loci within 0.2 cM from a divergent locus on the freshwater map (blue lines) and the F_1_ hybrid map (black lines).

We also observed stark differences between genetic maps. As expected, recombination in the hybrid map was completely suppressed within and immediately surrounding an inversion on chromosome 21 but this region recombined freely in the collinear BT map (length 4.3 cM, Figure 5). Conversely, on chromosome 1 the BT map showed bias in recombination toward chromosome ends expected on larger chromosomes. This bias was almost entirely absent in the hybrid map, with recombination occurring steadily throughout the chromosome. In both cases, the total map lengths of each chromosome were similar (chromosome 1: BT=129 cM, F_1_=137.5 cM; chromosome 21: BT=61.0 cM; F_1_=75.7 cM), indicating that these differences were due to differences in the distribution but not the number of crossovers.

### Much of the genome is tightly linked to loci under divergent selection

Genomic regions of greatest differentiation were compressed into proportionally smaller regions of the genetic maps (Figure 6, Table S2, Appendix 1). Some of the largest regions of differentiation we observed — including those surrounding *eda*, the major effect locus for lateral plate number (Colosimo *et al.* 2005), and the chromosome 21 inversion — had average F_ST_ values in excess of 0.4. Regions above this threshold spanned 8.1% of the physical genome (35.3 Mbp) but only 3.7% of the BT genetic map and 3.3% of the F_1_ hybrid genetic map (Figure S8). On chromosome 4, the large central regions of differentiation are similarly compressed on both genetic maps but this was not always the case. On chromosome 21, recombination was suppressed – and genomic differentiation compressed – only on the hererokaryotypic F_1_ map (Figure S1). A similar effect was seen across a large region of differentiation on chromosome 7 (Figure 6B) despite this chromosome being collinear between marine and freshwater stickleback genomes (Figure S1). On the physical map, chromosome 7 contained three peaks of strong differentiation between BT and RS spanning over 9 Mbp. All three peaks were clearly separated on the BT map, though they comprise a relatively smaller region than on the physical map, but collapsed to a single locus on the hybrid map (span: 0 cM, position 57.7 cM on the map).

Because our simulations suggested that population structure could interact with selection to maintain variation, but only with tight linkage (≤0.2 cM), we imputed the positions of RAD loci from the population genomic dataset onto the BT and hybrid genetic maps. Based on average genome-wide recombination rates, 0.2 cM equates to less than 50 kbp of sequence space (BT map: 36 kbp; F_1_ map: 44 kbp) and includes ~6% of sequenced loci. However, the clustering of divergent loci in regions of low recombination resulted in ~20% of all loci in the dataset fitting this linkage criterion (BT map: 19.5%; F_1_ map: 20.4%; Figure 6C,D; Figure S9). These tightly linked loci averaged over 200 kbp away from a divergent locus (BT map: 219 kbp; F_1_ map: 256 kbp) and included loci over 2 Mbp away (max, BT map: 2.3 Mbp; max, F_1_ map: 2.9 Mbp).

## Discussion

### Asymmetric population structure maintains asymmetries in patterns of linked genetic variation

The patterns of linked variation that we document here highlight not only the importance of selection but also the structure of the threespine stickleback metapopulation in maintaining genetic variation. Throughout the species range, the marine population is remarkably uniform phenotypically, which we now know is reflected genetically by minimal isolation-by-distance over thousands of kilometers (Bell and Foster 1994; Catchen *et al.* 2013a; Defaveri *et al.* 2013). This large population with few barriers to migration is contiguous with thousands of freshwater lake and stream populations that are more clearly isolated by geography. While the influence of freshwater populations on the evolution of the species was once unclear (Bell and Foster 1994), it is now evident that gene flow between freshwater and marine stickleback populations is common and may facilitate adaptation through the indirect sharing of alleles among freshwater populations (Schluter and Conte 2009).

Our findings suggest that the structuring of the freshwater habitat acts as a reservoir of genetic variation in the species, and that this effect is particularly potent in regions of the genome experiencing divergent selection. In genomic regions that are recombinationally distant to targets of divergent selection, gene flow among habitats homogenizes variation such that levels of genetic variation are similar (Figure 3) and differentiation is low (Figure 6) across habitats. At divergent loci themselves, selection has maintained a small number of haplotypes in each habitat and variation is partitioned almost entirely among habitats (Figure 2, Figure 3, Nelson and Cresko 2018). However, in the substantial fraction of the genome that is tightly linked to divergent loci, we hypothesize that selection has reduced the effective migration rate between alternative habitats such that the geographic structuring of the freshwater habitat becomes a significant factor affecting levels of genetic variation. Because migration among freshwater populations occurs primarily through the marine environment, divergent selection accentuates the effect of population structure in the freshwater habitat.

Forward simulations support this hypothesis. We found that under realistic migration rates (Caldera and Bolnick 2008; Berner *et al.* 2009) and selection coefficients (Barrett *et al.* 2008; Kitano *et al.* 2008), such asymmetric population subdivision *by itself* provides a simple explanation for the counterintuitive observation that smaller, isolated populations contain abundant linked variation (Figure 3, Figure 4). The degree to which population structure increased levels of diversity depended both on the amount of substructure and the length of time over which selection acted. While even minimal substructure eventually resulted in patterns of diversity mirroring those we observed in stickleback, this effect required a selection phase on the order of the expected coalescence time of the metapopulation. This follows from the fact that selection and population structure protect variants from fixation or loss at adaptive and linked loci, respectively. Our findings generally agree with expectations from Slatkin’s (1991) model, where limited migration among demes increases the expected coalescence times of chromosomes sampled from a metapopulation. Nearby an alternatively adaptive locus, selection accentuates this effect by reducing effective migration rates among demes of alternative habitats.

Our empirical and simulated results, in combination with previous findings in threespine stickleback (Reimchen 1994; Kaeuffer *et al.* 2012; Roesti *et al.* 2014), have important implications for the long-term evolution of the species. Freshwater populations are ecologically diverse (Bell and Foster 1994; Mckinnon and Rundle 2002; Kaeuffer *et al.* 2012; Leaver and Reimchen 2012; Reimchen *et al.* 2013) and the relative impacts of parallel versus non-parallel selection pressures among freshwater populations is actively debated (Kaeuffer *et al.* 2012; Stuart *et al.* 2017). While it is possible that multifarious selection pressures also contribute to the maintenance of more genetic diversity on chromosomes from freshwater fish, our model does not require additional selection to maintain linked variation. Furthermore, the freshwater pond populations studied here are ecologically similar and geographically proximal (Cresko *et al.* 2004), so it is likely that parallel selection on a common pool of variation is largely responsible for the patterns of variation we observe. Put another way, we find no need to invoke more complex selection regimes in order to explain the patterns of variation we observe. Also studying ecologically similar freshwater populations, Roesti et al (2014) found that parallel adaptation led to increases in F_ST_ among freshwater populations near peaks of marine-freshwater divergence, and inferred that sweeps of common SNPs on different genetic backgrounds led to this pattern. We have shown these linked genomic regions to be not only the most heavily partitioned but also the most diverse in the genome (Figure 3F, Figure S6). We find it likely that selection and gene flow happening over shorter evolutionary timescales (tens to thousands of years), like those investigated by Roesti et al. (2014), have occurred throughout the evolutionary history of this species and have long-lasting, collective effects on patterns of genomic variation over the course of millions of years.

### A variable recombination landscape increases the genomic reach of selection and simplifies the architecture of divergence

In addition to the consequences of asymmetric population structure in maintaining genetic variation unevenly across geography, the recombination landscape also can have profound effects on heterogenous genome-wide patterns of genetic variation. Linkage to or among selected sites is now thought to affect many, if not most, variable sites in the genome (Schrider and Kern 2017; Kern and Hahn 2018), and suppression of recombination among the genomes of diverging populations and species appears in large part to determine patterns of genetic differentiation and divergence (Burri *et al.* 2015; Samuk *et al.* 2017; Vijay *et al.* 2017; Stankowski *et al.* 2018). In the threespine stickleback, genetic differentiation among phenotypically divergent populations is known to accumulate in regions of low recombination (Roesti *et al.* 2013; Marques *et al.* 2016; Samuk *et al.* 2017). Our work adds an important outcome of this phenomenon: low recombination adjacent to targets of divergent selection extends selection’s reach into linked regions, maintaining genetic variation and structuring it among chromosomes with divergent evolutionary histories.

The accumulation of genetic diversity adjacent to targets of selection required tight linkage in our simulations, potentially limiting the relevance of this model in explaining patterns of genetic variation. However, because adaptive divergence has occurred principally in regions of low recombination, a substantial fraction of the genome is likely near enough (~0.2 cM) to divergent loci for our model to explain the accumulation of variation on freshwater chromosomes. The absolute genetic distance through which divergent selection can influence linked variation is proportional to the strength of selection (e.g. Figure 4) and inversely proportional to the migration rate (e.g. Figure S10, Charlesworth *et al.* 1997). While we did not fully explore either parameter space, our chosen values are congruent with empirical estimates from stickleback. Estimates of migration rates vary widely across studies of stickleback, from nearly zero to rates that could swamp divergent selection (Caldera and Bolnick 2008; Berner *et al.* 2009). Estimated selection coefficients on lateral plate armor are consistently above 0.2 (Barrett *et al.* 2008; Kitano *et al.* 2008), and the total effect of selection on a locus depends on the total number of nearby selected sites (i.e. the ‘selection density’: Aeschbacher *et al.* 2017). Given the rate of adaptation observed in recently colonized freshwater habitats (Lescak *et al.* 2015; Bassham *et al.* 2018) combined with generally low recombination rates, the effect of selection may at times extend across entire chromosomes.

Our results also provide general insight into the variability and evolution of the recombination landscape. First, genomic divergence that occurs in the centers of the largest chromosomes (Hohenlohe *et al.* 2010; Jones *et al.* 2012; Bassham *et al.* 2018; Nelson and Cresko 2018) occurs within regions of consistently low recombination across maps from different genetic backgrounds(Hohenlohe *et al.* 2012; Roesti *et al.* 2013; Samuk *et al.* 2017). The negative association between divergence and recombination rate is a common finding across systems (Nachman 2002; Carneiro *et al.* 2009; Geraldes *et al.* 2011; Cutter and Payseur 2013; Burri *et al.* 2015; Aeschbacher *et al.* 2017), and can result from both negative (Burri *et al.* 2015; Stankowski *et al.* 2018) and positive selection (Aeschbacher *et al.* 2017; Samuk *et al.* 2017) between allopatric populations (Burri *et al.* 2015) and those currently exchanging genes (Carneiro *et al.* 2009; Aeschbacher *et al.* 2017). Our results further demonstrate that stickleback are no exception to this rule and that divergent selection in the face of (historical) gene flow has generated this relationship at both at the local (i.e. specific genomic regions, Figure 6) and genome-wide scales (Table S2, Figure S8). Second, although these results generally hold using both intra- and inter-population genetic maps, we find that recombination rates in regions of divergence were often lowest in the F_1_ hybrid map (Figure 5; Figure 6B; Figure S8). While this was expected within chromosomal inversions, where the alternative homozygotes demonstrated steady recombination throughout the inverted region, this was also the case on chromosome 7 despite the apparent lack of any large scale, simple structural variation (Figures 2 and 4) although clusters of smaller structural variants that could not be detected in our maps could also reduce recombination. On the hybrid map, this entire region collapsed to a region inherited essentially as a single Mendelian locus.

The specific reductions in recombination on the hybrid map are suggestive of the evolution of the recombination landscape itself and, if true, could have profound implications for adaptation and genomic divergence in stickleback. For example, inversions are advantageous when recombination between multiple linked alleles that contribute to fitness (either in an additive or epistatic fashion) is maladaptive (Kirkpatrick and Barton 2006). The three previously identified inversions on chromosomes 1, 11 and 21 (Jones *et al.* 2012) are associated with divergence between the freshwater and marine habitats and occur in regions of the genome that readily recombine in chromosomal homozygotes (Figure 4, Figure S1, Figure S2). Had inversions not evolved, gene flow and recombination among marine and freshwater populations may have been strong enough to prevent adaptive divergence in these genomic regions (Lenormand 2002; Yeaman and Whitlock 2011; Aeschbacher *et al.* 2017).

In contrast, our work and that of others (Roesti *et al.* 2013; Glazer *et al.* 2015) has shown that megabase pair-scale inversions have not evolved across the largest regions of divergence in the genome. In these regions, low recombination rates — which may themselves have evolved adaptively or been ancestral platforms for adaptive divergence — combined with strong selection has been effective at maintaining allelic combinations across megabase pairs of genomic space. These are also regions of exceptional sequence divergence between marine and freshwater chromosomes (Table S2; Samuk *et al.* 2017; Nelson and Cresko 2018). Sequence divergence alone may limit double-strand break resolution as crossovers in these regions (Modrich and Lahue 1996; Opperman *et al.* 2004; Li *et al.* 2006), biasing crossovers toward regions of lower sequence divergence. Explicit tests of this hypothesis, for example with F_3_ mapping families where the effects of marine-freshwater hetero- and homozygosity in specific genomic regions can be directly assessed, will be a productive avenue of research.

The coincidence of reduced recombination and genomic divergence may help explain the well documented pattern of repeatable and rapid adaptive divergence in stickleback (Bell and Foster 1994; Lescak *et al.* 2015) that is largely the result of the reuse of standing genetic variation (Colosimo *et al.* 2005; Schluter and Conte 2009; Deagle *et al.* 2012; Terekhanova *et al.* 2014; Roesti *et al.* 2015; Marques *et al.* 2016; Bassham *et al.* 2018). Because freshwater populations are typically founded by marine stickleback, the sources of standing genetic variation are low frequency variants in the marine population that nearly always exist in a heterozygous state with a marine genome (Bassham *et al.* 2018). Our results suggest that the recombination landscape may therefore facilitate the maintenance of freshwater haplotypes during their transit through the marine environment by reducing recombination even in collinear genomic regions. Multi-megabase haplotypes — potentially containing many alleles contributing to local adaptation — then have a higher probability of being selected in concert when a new freshwater population is established.

## Conclusions

Divergent natural selection is a powerful force for the maintenance of genetic variation in nature. When local adaptation plays out on variable geographic and recombinational landscapes, we find that the effect of selection on genetic variation are amplified and shape patterns of linked variation in unexpected ways. Here we document using simulation and empirical studies that asymmetric population subdivision among habitats in stickleback leads to an overall greater maintenance of diversity in freshwater as compared to the larger ancestral marine population. Furthermore, we hypothesize that the stickleback recombinational landscape is the product of repeated adaptive evolution that transforms a large genomic architecture of marine-freshwater divergence into a much more simplified genetic architecture. If true, two consequences are the efficient maintenance of adaptive divergence in the face of gene flow, and widespread linked selection that eliminates genetic variation in the panmictic marine population while maintaining genetic diversity in the freshwater habitat. The interaction of selection, recombination, and population structure has turned small, isolated freshwater stickleback populations into primary reservoirs of standing genetic variation that may potentiate future evolutionary change. More broadly, our results underscore how integrated studies of evolutionary and genetic processes can yield exciting, unexpected patterns and deeper understanding when we jointly consider the processes and their interactions with one another.

## Author Contributions

TCN, JMC, and WAC conceived of and designed the study. TCN, JGC, CMI, and JMC performed mapping crosses and prepared sequencing libraries. TCN, JGC, and JMC performed linkage mapping. TCN performed population genomic analyses and conducted population genetic simulations. TCN and WAC wrote the paper.

## Acknowledgements

We thank Patrick Phillips, John Postlethwait, Kirstin Sterner, Matt Streisfeld, and members of the Cresko Laboratory for their guidance and advice. Thanks go to Sean Stankowski, Nadia Singh, Peter Ralph, Lila Fishman, Madeline Chase, Emily Beck, Kristin Alligood and the evolutionary genetics group at the University of Montana for advice for useful discussions throughout the development of this project. And thanks to Katie Peichel, David Begun and three anonymous reviewers for their comments on and improvements to this manuscript. We acknowledge National Science Foundation awards NSF DEB 1501423 (W.A.C. and T.C.N.), NSF DEB 0919090 (W.A.C.), and National Institutes of Health award NIH T32GM007413 (T.C.N.).

## Supplementary Tables and Figures

**Table S1.**
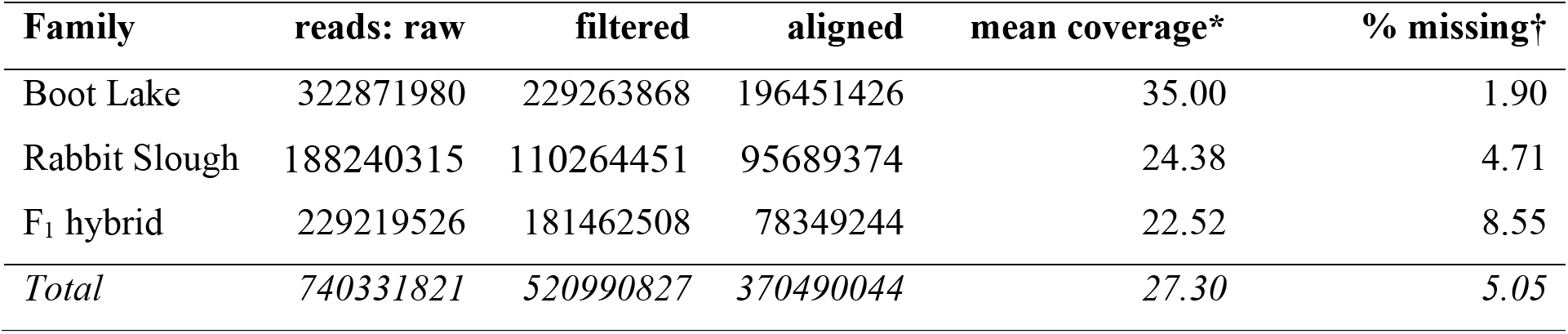
Sequencing statistics of libraries for mapping families in the main text. *Coverage is the mean sequencing depth per RAD marker across the 94 progeny with highest coverage. These progeny were used to estimate recombination rates in the main text. †Missing data calculated as the average percent of RAD markers per progeny with zero coverage.

| Family | reads: raw | filtered | aligned | mean coverage* | % missing† |
| --- | --- | --- | --- | --- | --- |
| Boot Lake | 322871980 | 229263868 | 196451426 | 35.00 | 1.90 |
| Rabbit Slough | 188240315 | 110264451 | 95689374 | 24.38 | 4.71 |
| F<sub>1</sub> hybrid | 229219526 | 181462508 | 78349244 | 22.52 | 8.55 |
| *Total* | *740331821* | *520990827* | *370490044* | *27.30* | *5.05* |

**Table S2.** Genome-wide correlations between recombination rate and genetic diversity and divergence. Statistics were calculated as means of 1000-kb non-overlapping bins. π_RS_: sequence diversity in the marine RS population. π_FW_: sequence diversity in the combined freshwater populations. Correlation coefficients are given as Spearman’s ρ. Characters in parentheses indicate significant correlations at p < 0.01: *linear model; ρ = Spearman’s ρ, τ = Kendall’s τ.

| map | $\pi_{RS}$ | $\pi_{FW}$ | $\pi$ | $d_{XY}$ | $F_{ST}$ |
| --- | --- | --- | --- | --- | --- |
| BT | 0.450 (*,$\rho$,$\tau$) | 0.349 (*,$\rho$,$\tau$) | 0.261 (*,$\rho$,$\tau$) | 0.154 ($\rho$,$\tau$) | -0.431 (*,$\rho$,$\tau$) |
| RS | 0.489 (*,$\rho$,$\tau$) | 0.223 (*,$\rho$,$\tau$) | 0.225 (*,$\rho$,$\tau$) | 0.146 ($\rho$,$\tau$) | -0.365 ($\rho$,$\tau$) |
| F<sub>1</sub> | 0.483 (*,$\rho$,$\tau$) | 0.243 (*,$\rho$,$\tau$) | 0.146 ($\rho$,$\tau$) | 0.053 | -0.406 (*,$\rho$,$\tau$) |

**Figure S4.**
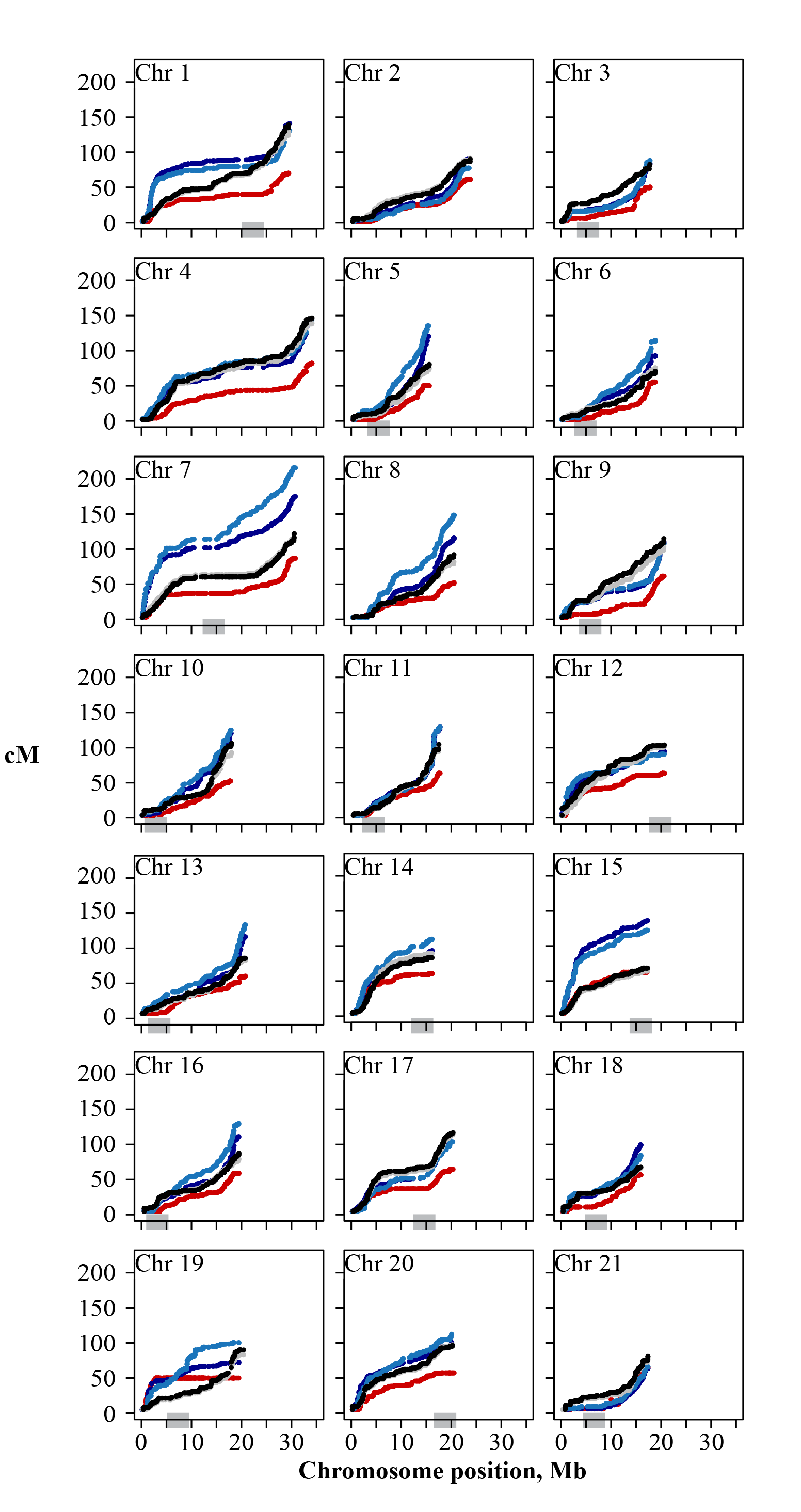
Genetic maps for all 21 threespine stickleback chromosomes. Genetic position for each RAD marker is plotted against aligned physical position on the threespine stickleback reference genome. Boot Lake: dark blue; Boot Lake, subset of 94 progeny: blue; Rabbit Slough: red; F_1_ hybrid: gray; F_1_ hybrid, subset of 94 progeny: black. Gray boxes indicate the approximate positions of centromeres (where known) from Cech and Peichel (2015) and Sardell et al (2018).

**Figure S2.**
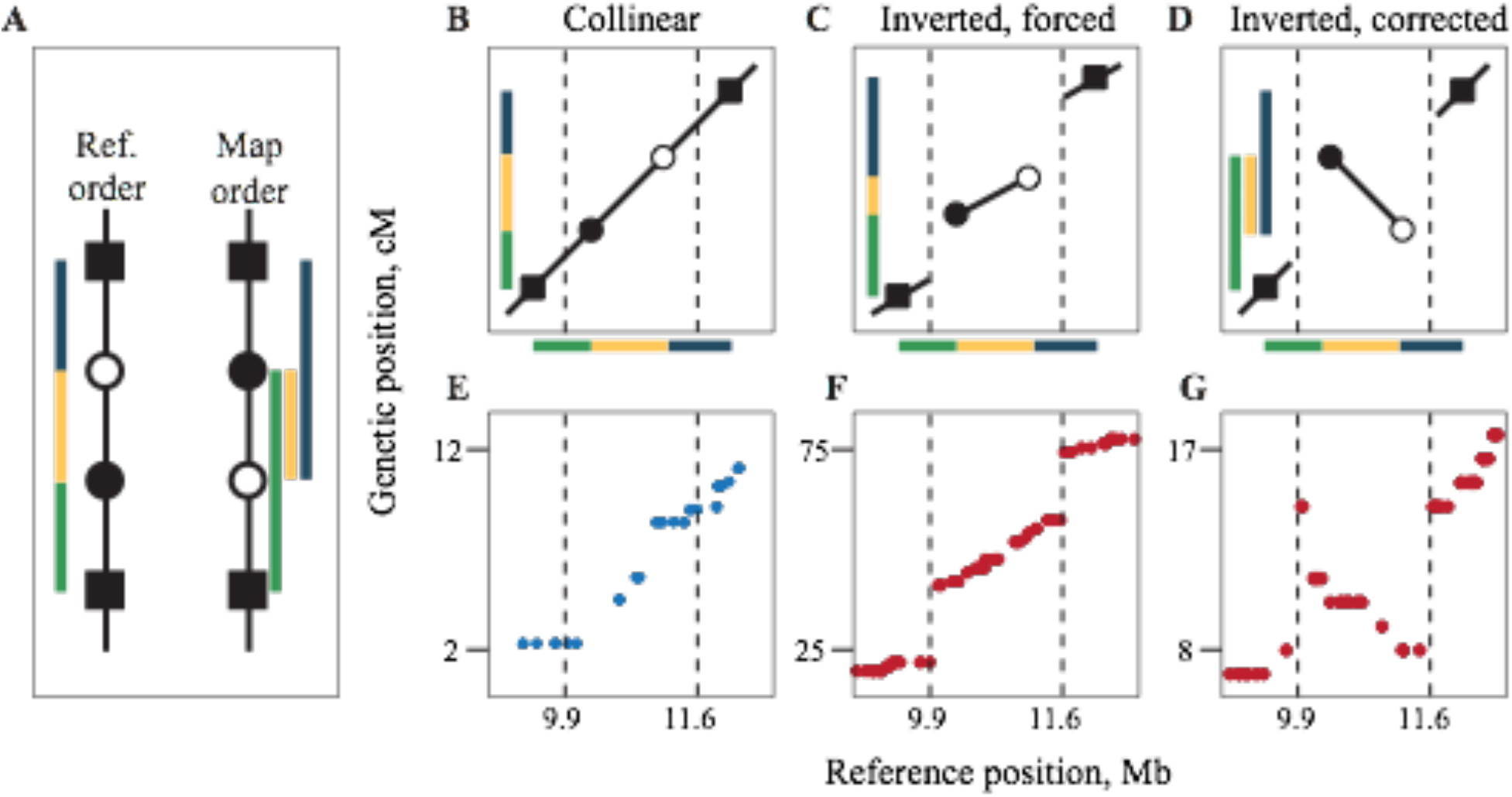
Reference-guided genetic map construction allows identification and correction of chromosomal inversions. A: Inversion polymorphisms create discrepancies between the reference physical genome and a genetic map. Colored bars indicate true map distances between genetic markers inside and on either side of the inversion. B: When the physical and genetic maps are collinear, map distances agree with the reference. C: An inverted genetic map forced into reference order results in a disjunct genetic map because crossover frequencies between markers inside and outside the inversion are inferred correctly. Note that the green and blue bars reflect the true distances between the markers involved but they are in the wrong order. D: Reversing the order of markers within the inversion preserved the relative map distances and reduced the overall map length. E-G: Genetic maps from the BT fish (E, collinear with reference) and the RS fish (F and G, inverted relative to reference) demonstrate this correction using empirical data.

**Figure S3.**
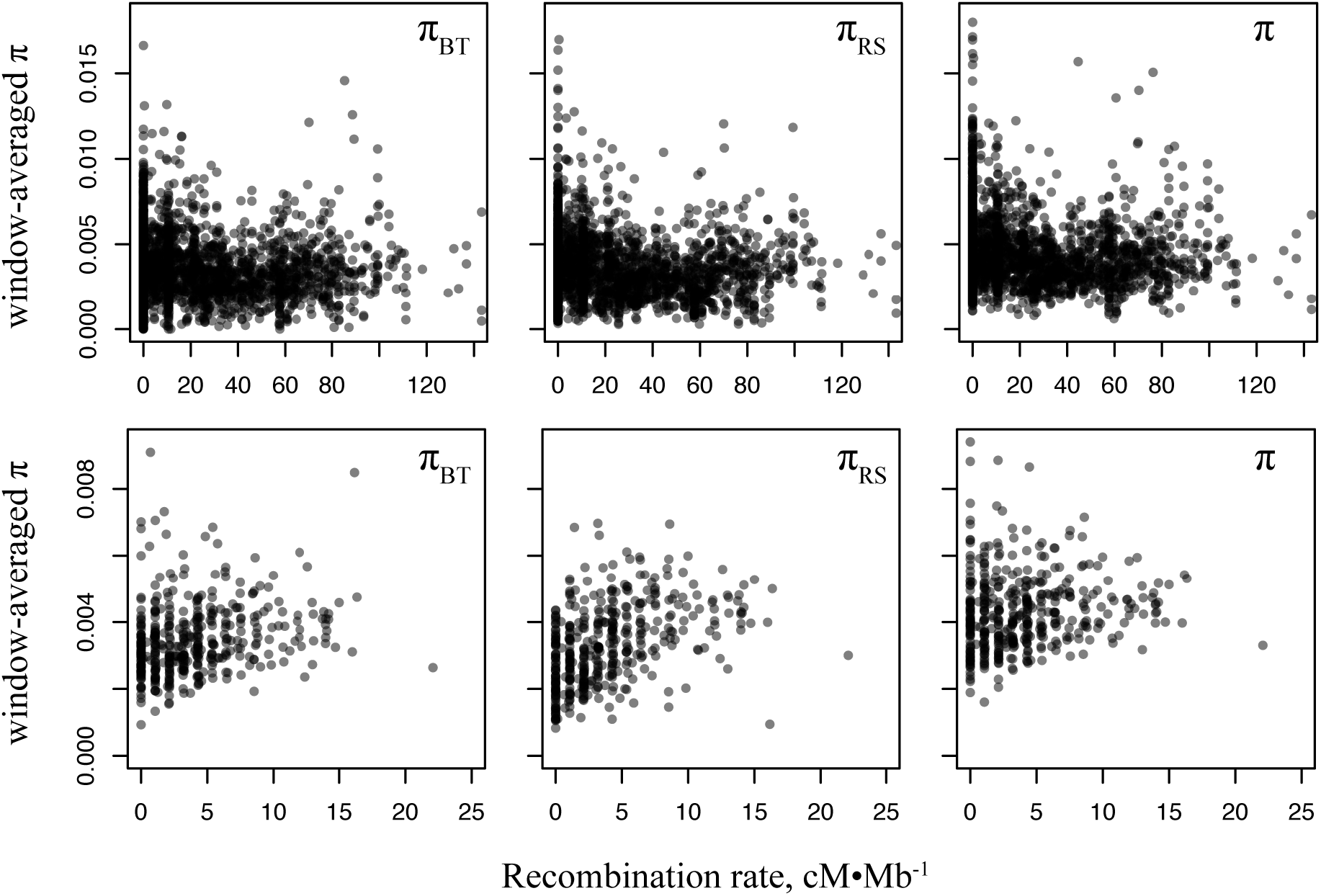
Window-based estimates of recombination rate (cM/Mb) and sequence diversity using (top) 100-kb non-overlapping windows and (bottom) 1-Mb nonoverlapping windows. Note the difference in axis scale with different window sizes.

**Figure S4.**
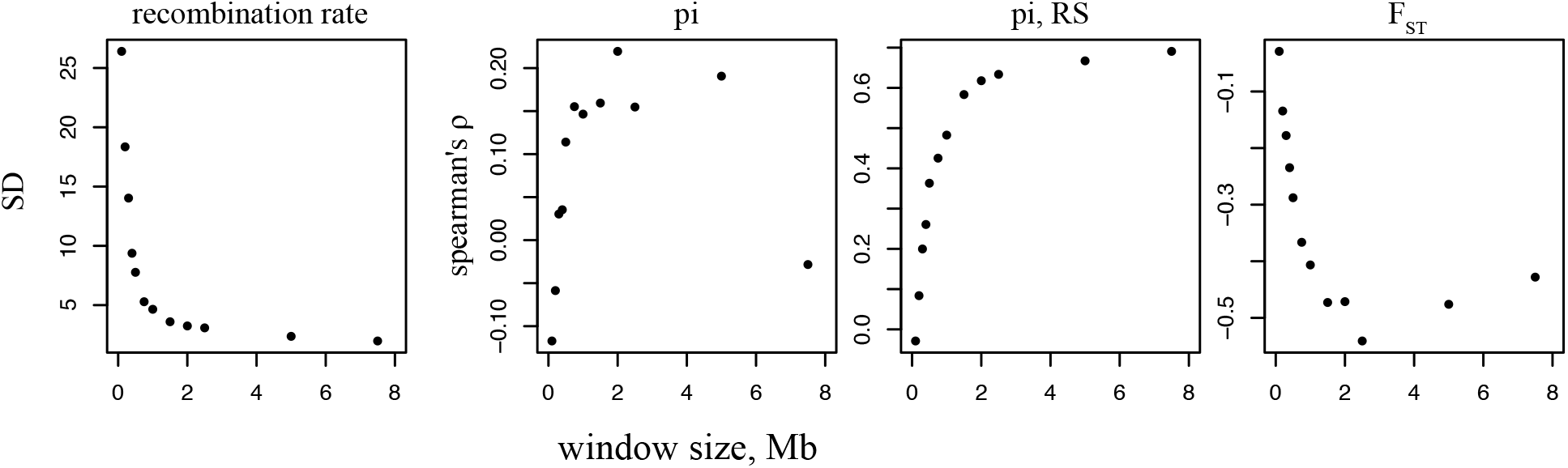
Variation in recombination rate and covariation with population genetic statistics at different genomic window sizes. All points are genome-wide summaries of recombination rate (cM/Mb) and correlation (as Spearman’s ρ) with population genetic variation and differentiation at a given window size using non-overlapping genomic windows. SD: standard deviation; π: sequence diversity; F_ST_: differentiation between Rabbit Slough and Boot Lake populations.

**Figure S5.**
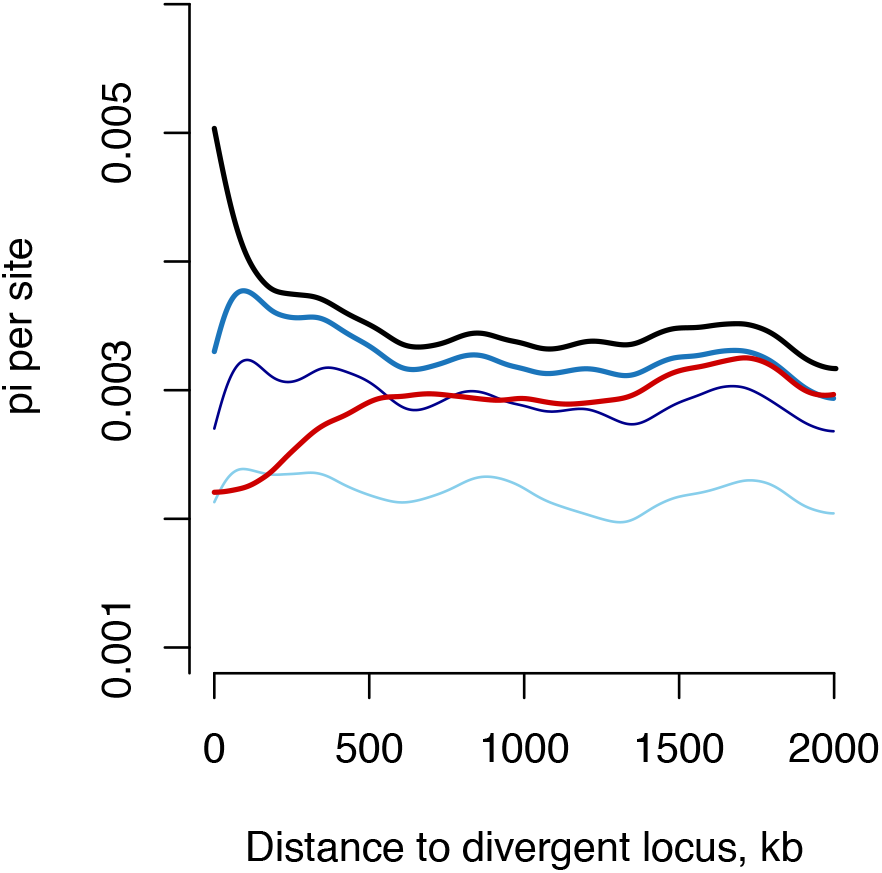
Sequence diversity as a function of distance from divergent loci in all populations. All genomes: black; Rabbit Slough: red; combined freshwater: middle blue; Boot Lake: dark blue; Bear Paw Lake: light blue.

**Figure S6.**
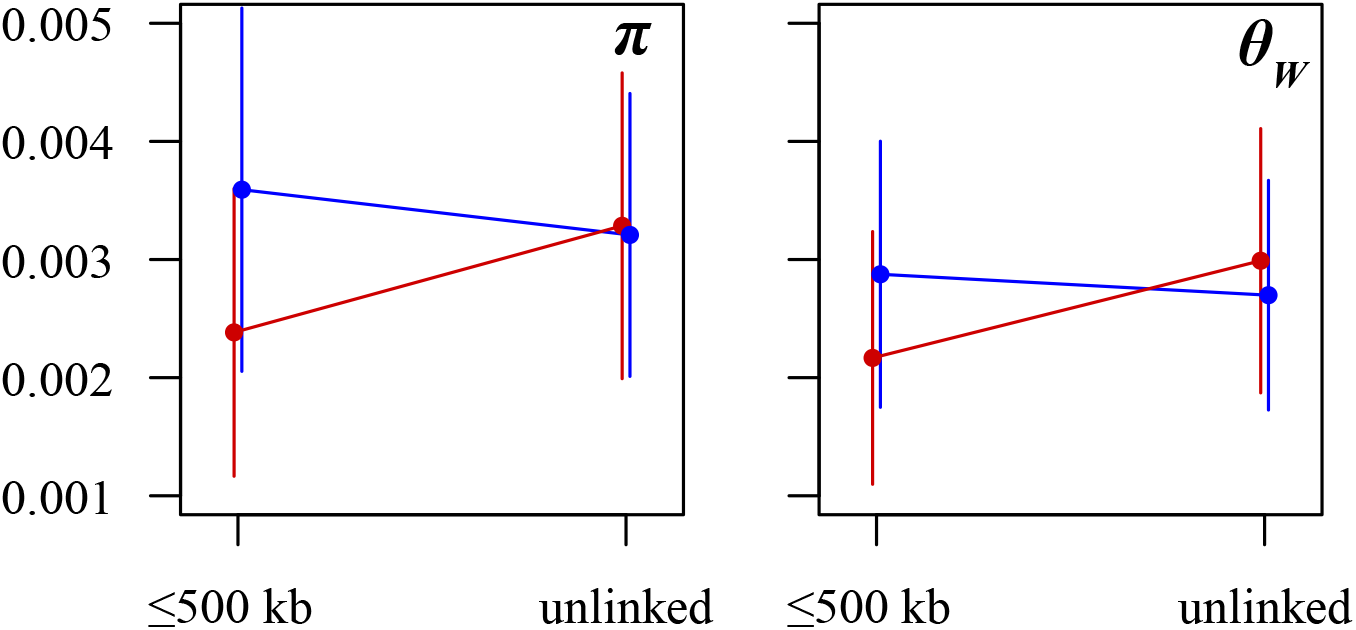
Interactions between genetic diversity and distance from divergent loci for both π and θ_W_. Rabbit Slough chromosomes: red; combined freshwater chromosomes: blue. All comparisons are significant (Tukey’s HSD: p ≤ 0.01) except Rabbit Slough and freshwater π at unlinked loci.

**Figure S7.**
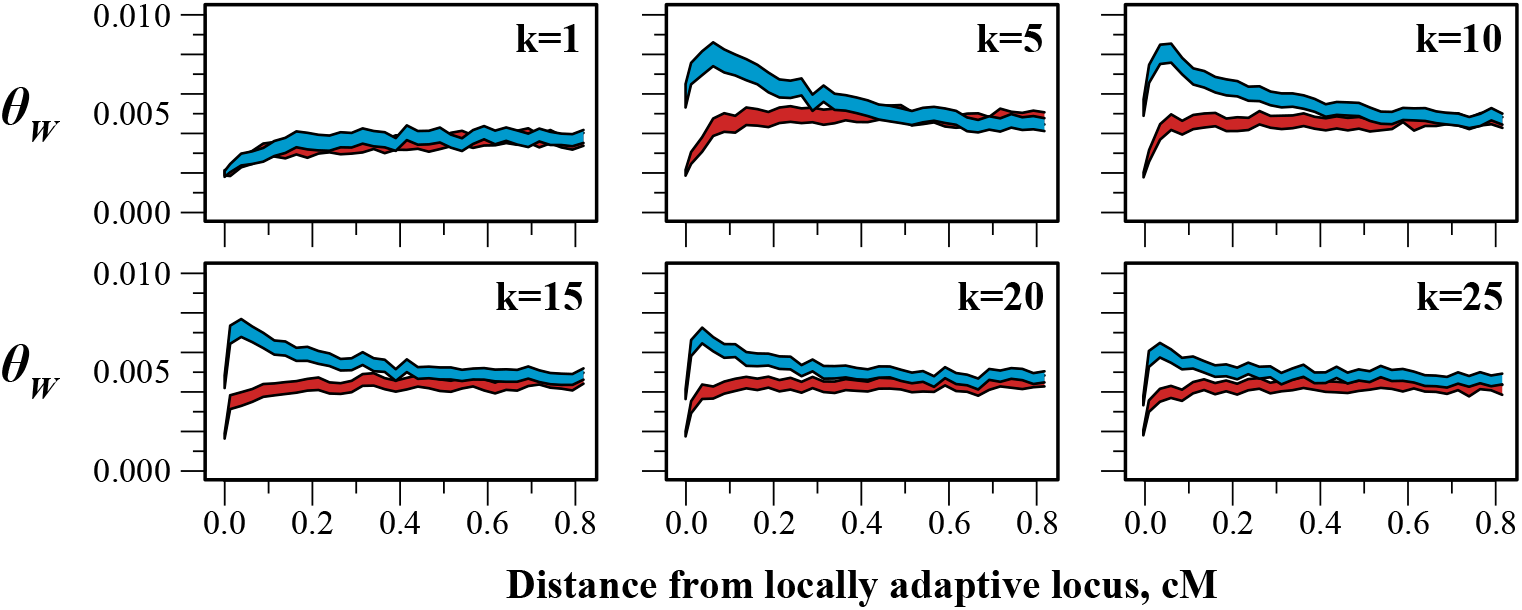
Effects of population subdivision linked genetic variation. All simulations were run as in the main text, Figure 4: s = 0.20; m = 1 migrant/generation; 20N_H_ generations of selection.

**Figure S8.**
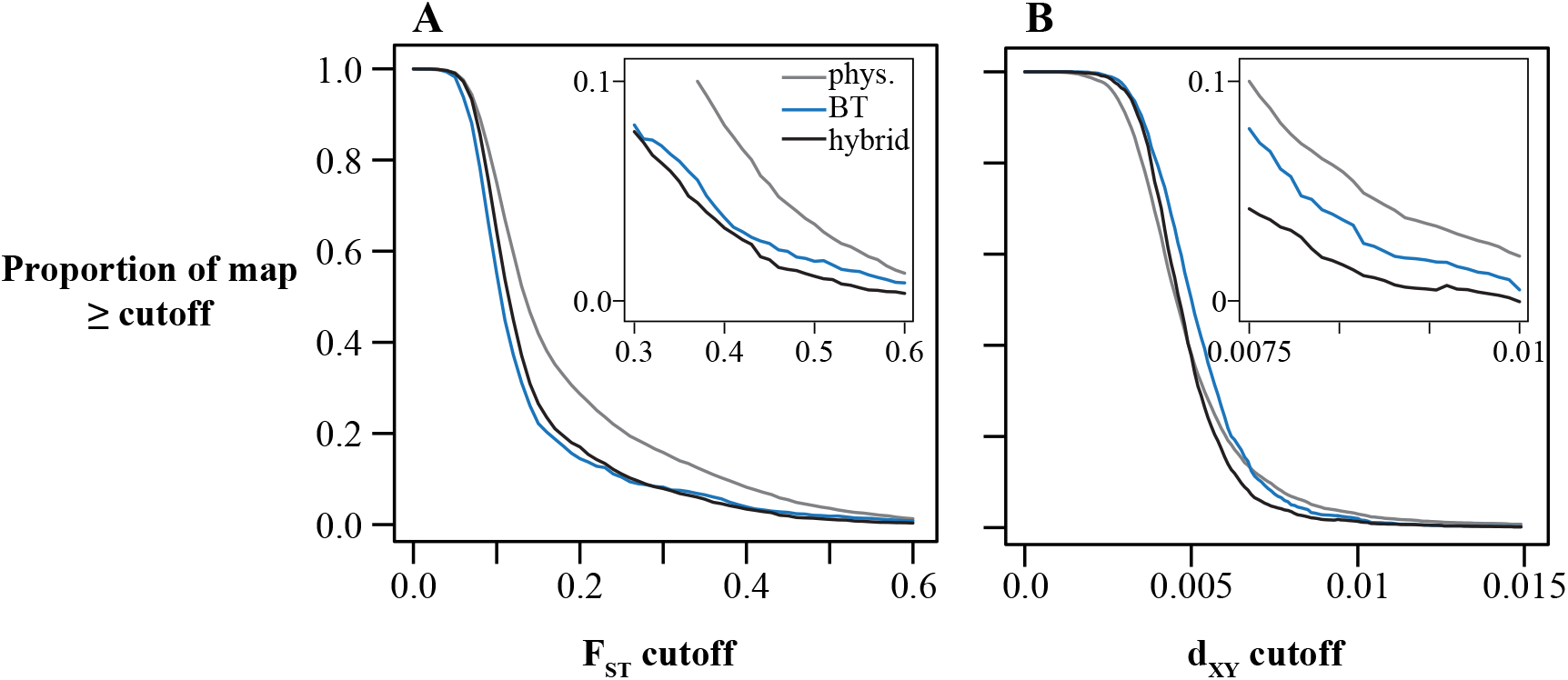
Genetic divergence comprises a smaller proportion of the genetic map than the physical map. Each line represents the proportion of the overall map lengths (Mb or cM) taken up by windows of genetic divergence (F_ST_ or d_XY_) greater than or equal to a given value. Gray: physical map; blue: Boot Lake genetic map; black: F_1_ hybrid genetic map.

**Figure S9.**
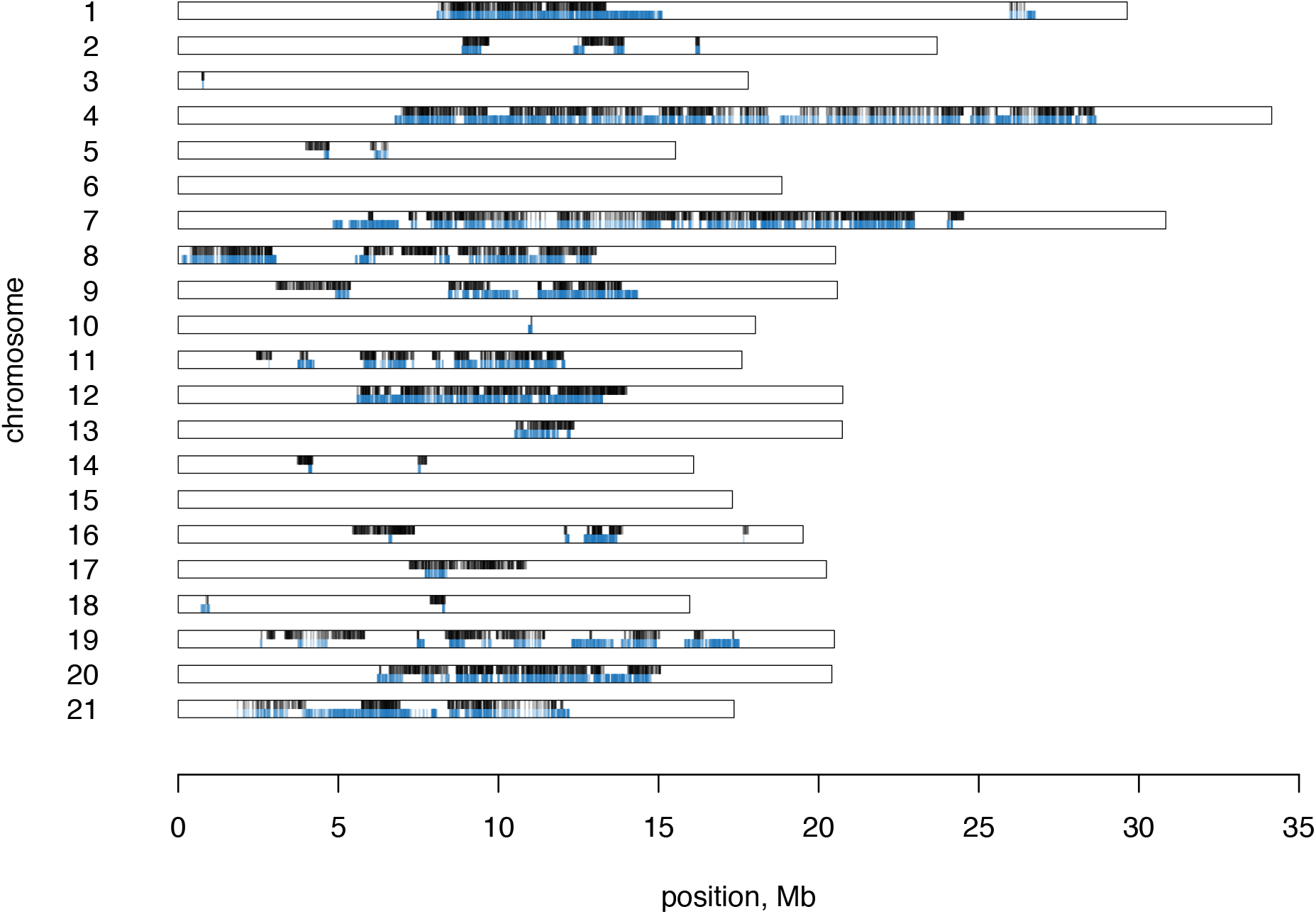
Locations of all RAD loci in the population genomics dataset with imputed genetic positions within 0.2 cM of a divergent locus. F_1_ hybrid genetic map: black; Boot Lake genetic map: blue.

**Figure S10.**
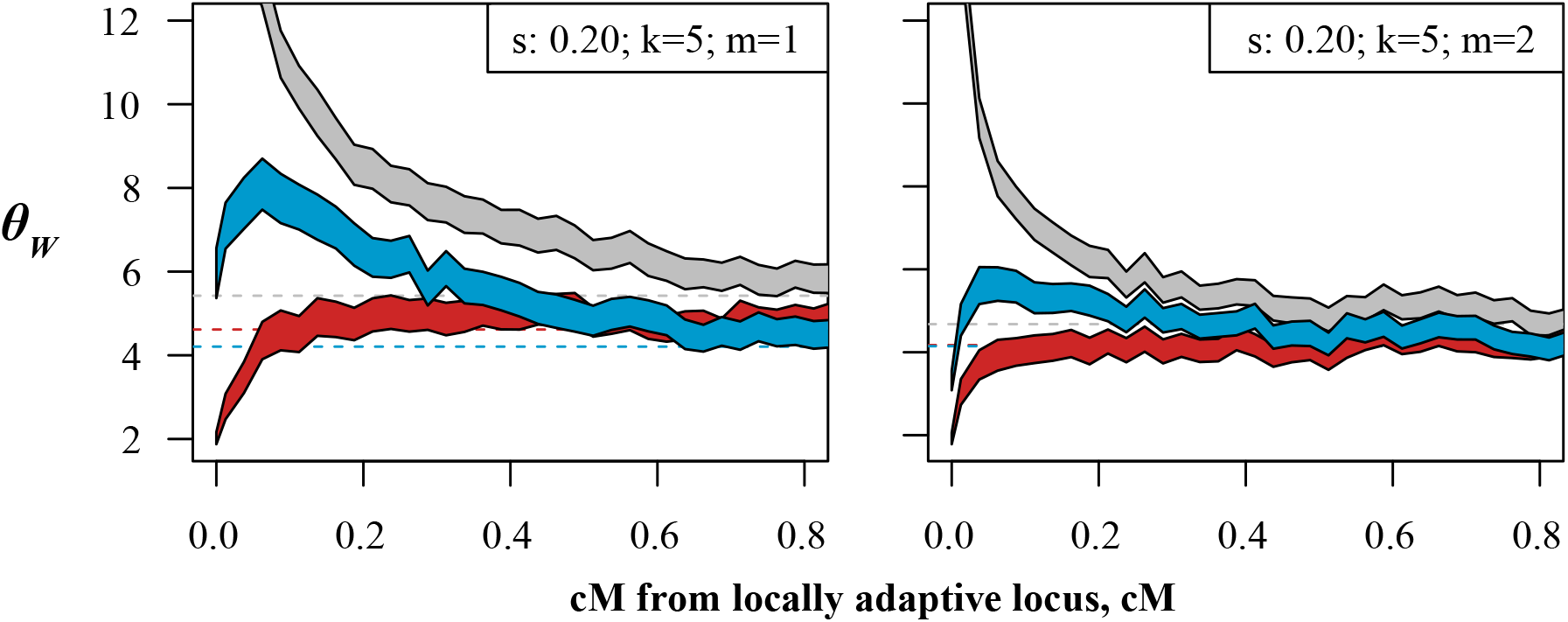
Effect of migration rate on linked genetic variation. All simulations were run for 20N_H_ generations of selection.

## Appendix 1. Physical-to-genetic map scans for all chromosomes

Maps plotted as per Figure 6 in the main text. Grayscale reflects window-averaged F_ST_. Gold lines are positions of divergent loci (Nelson and Cresko 2018).

**Figure 1:**
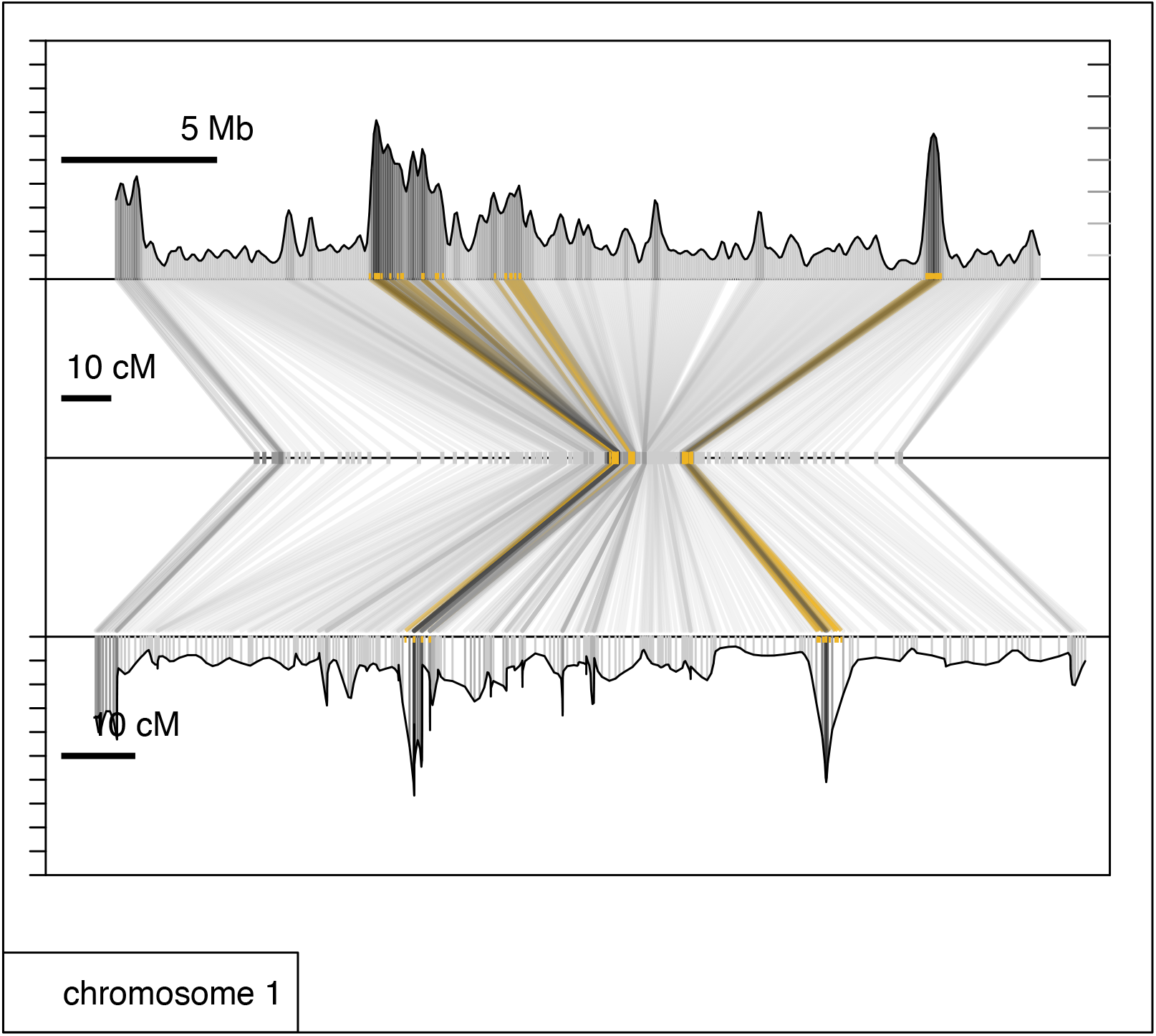
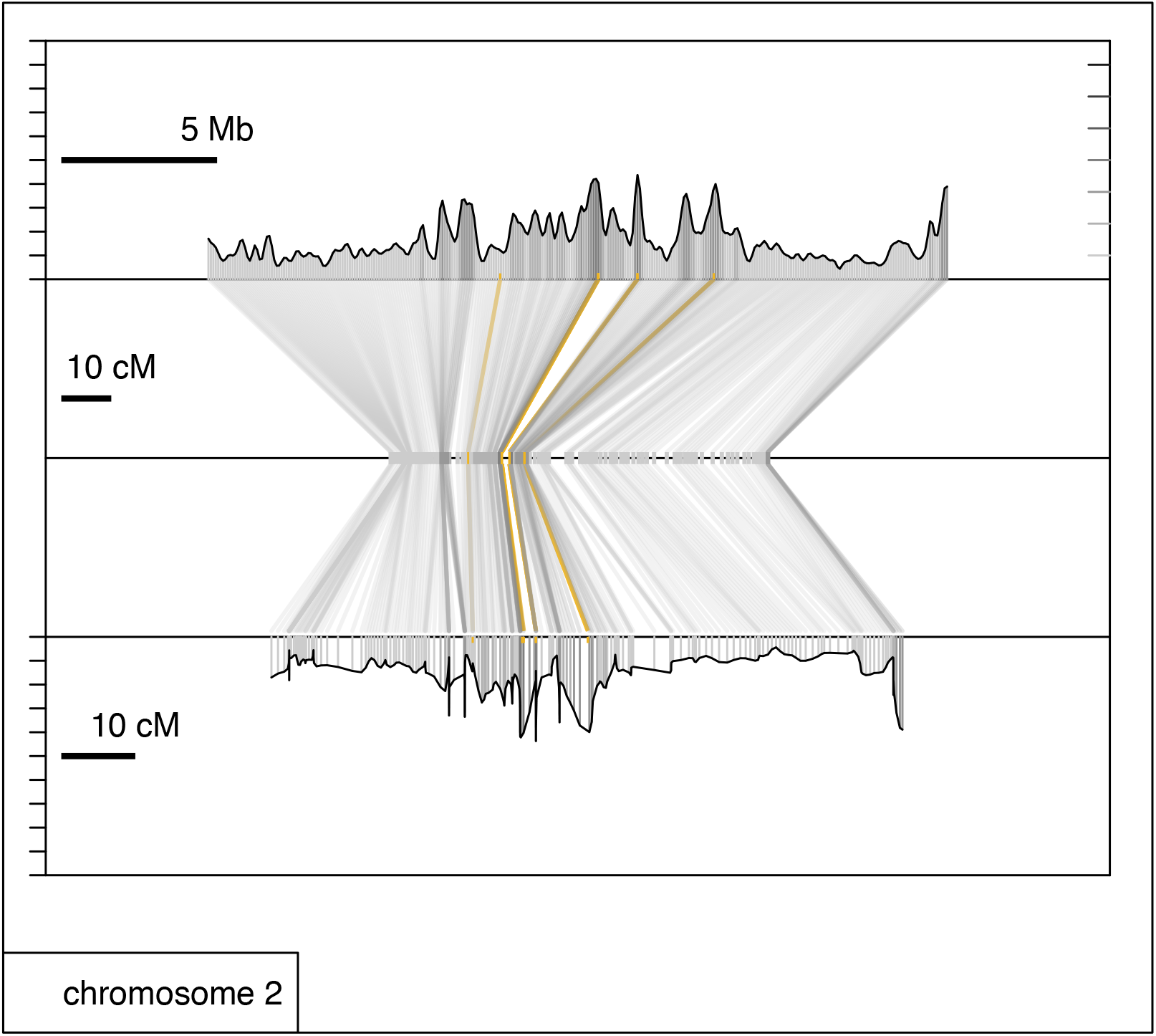
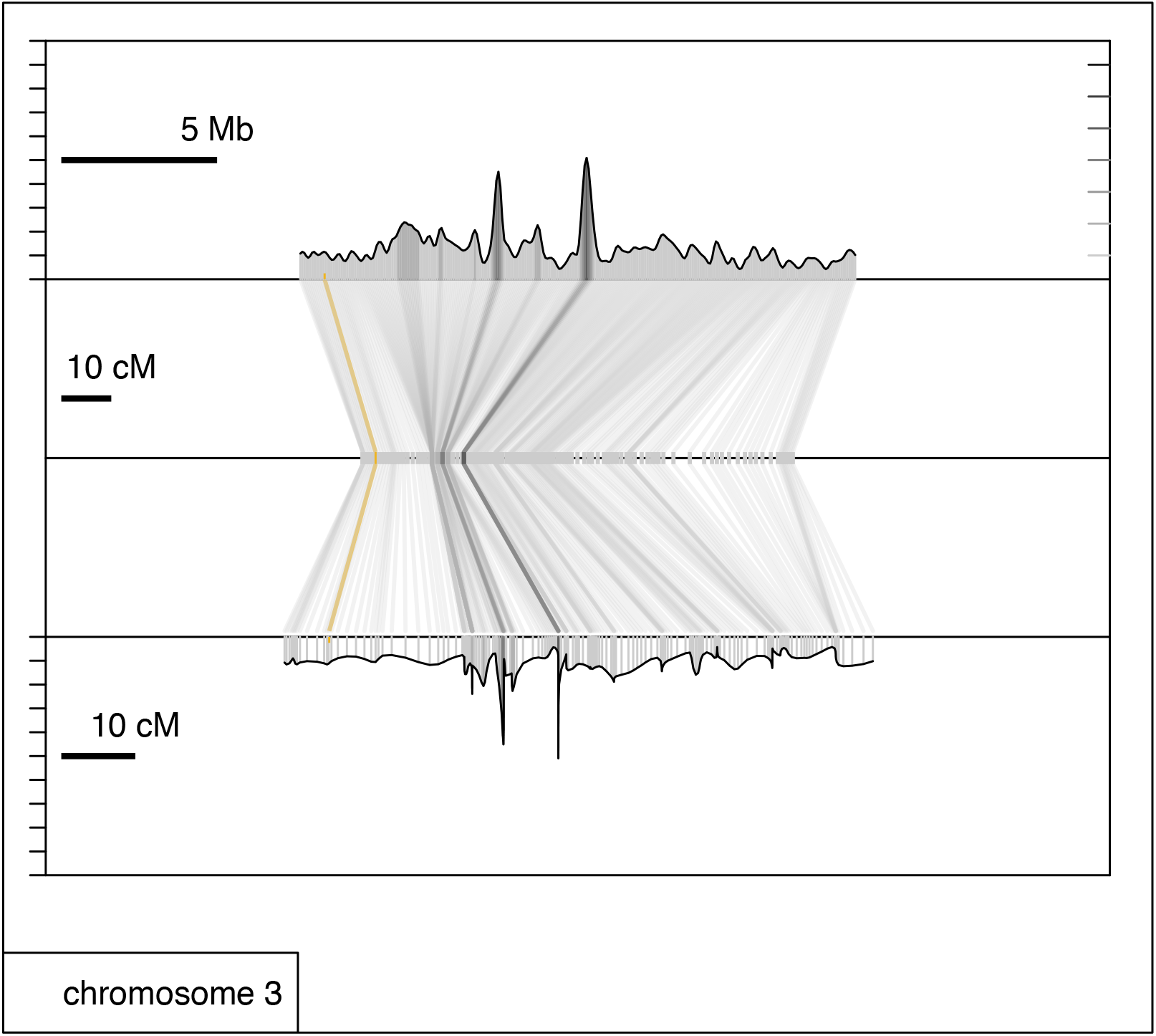
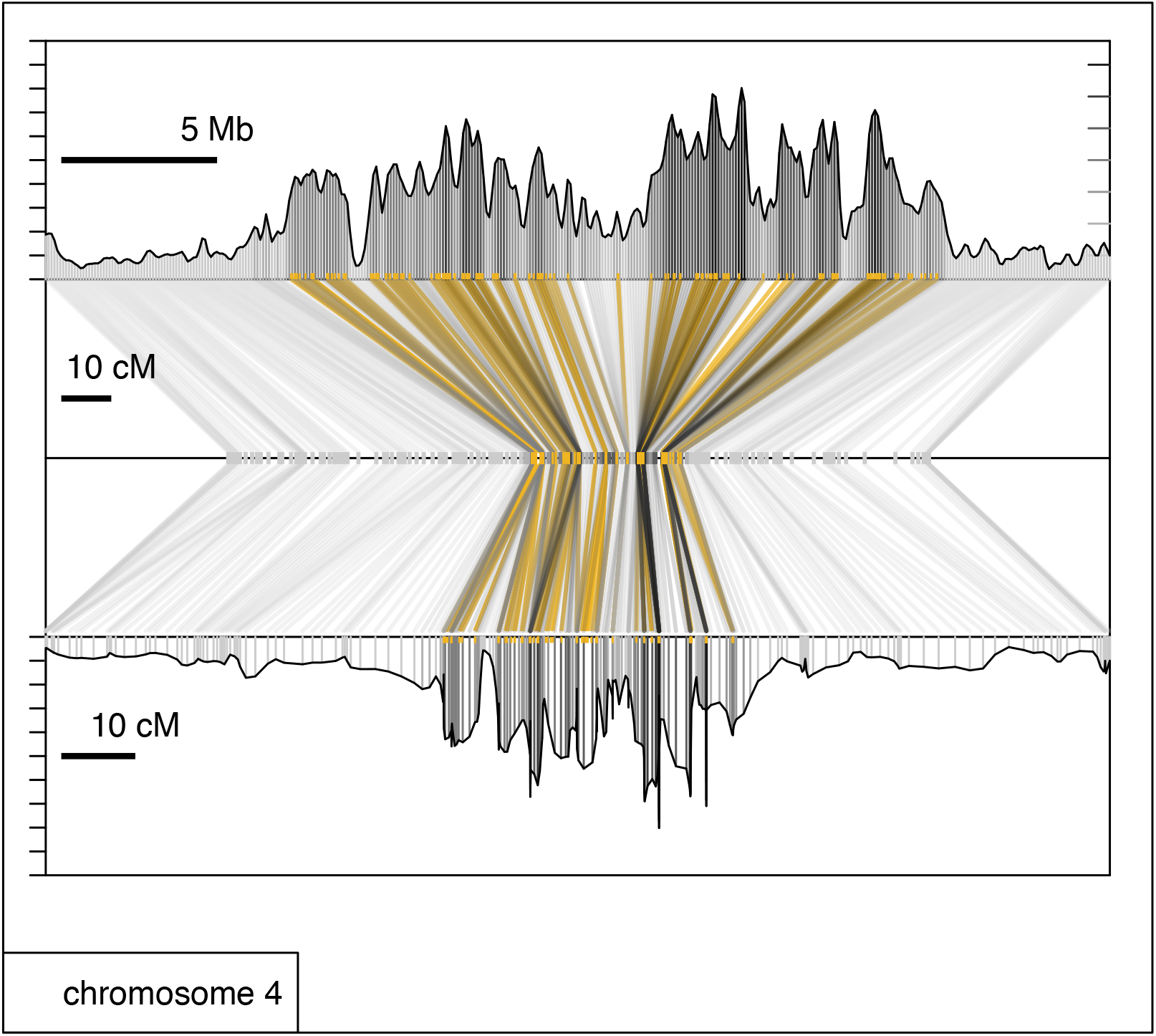
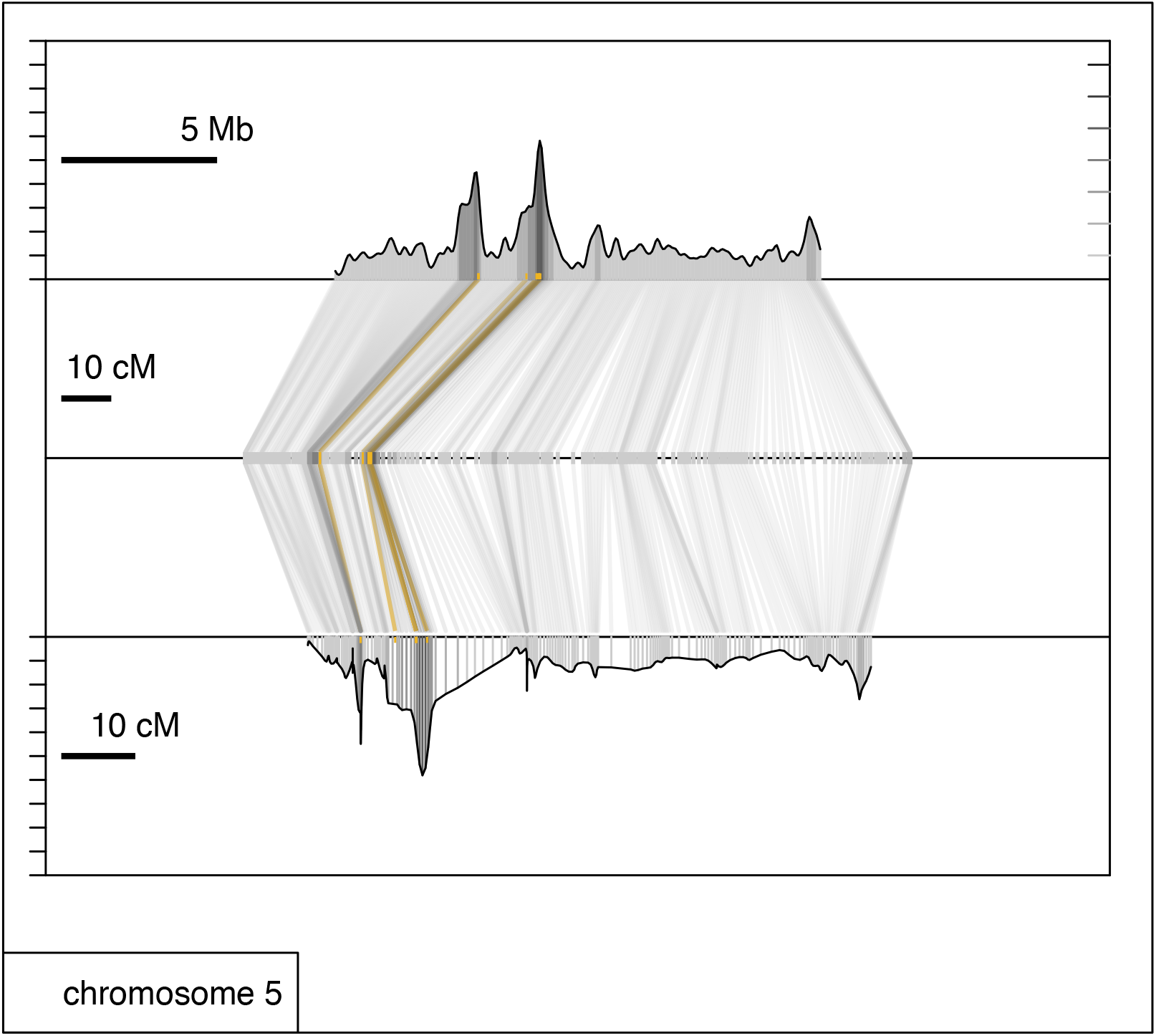
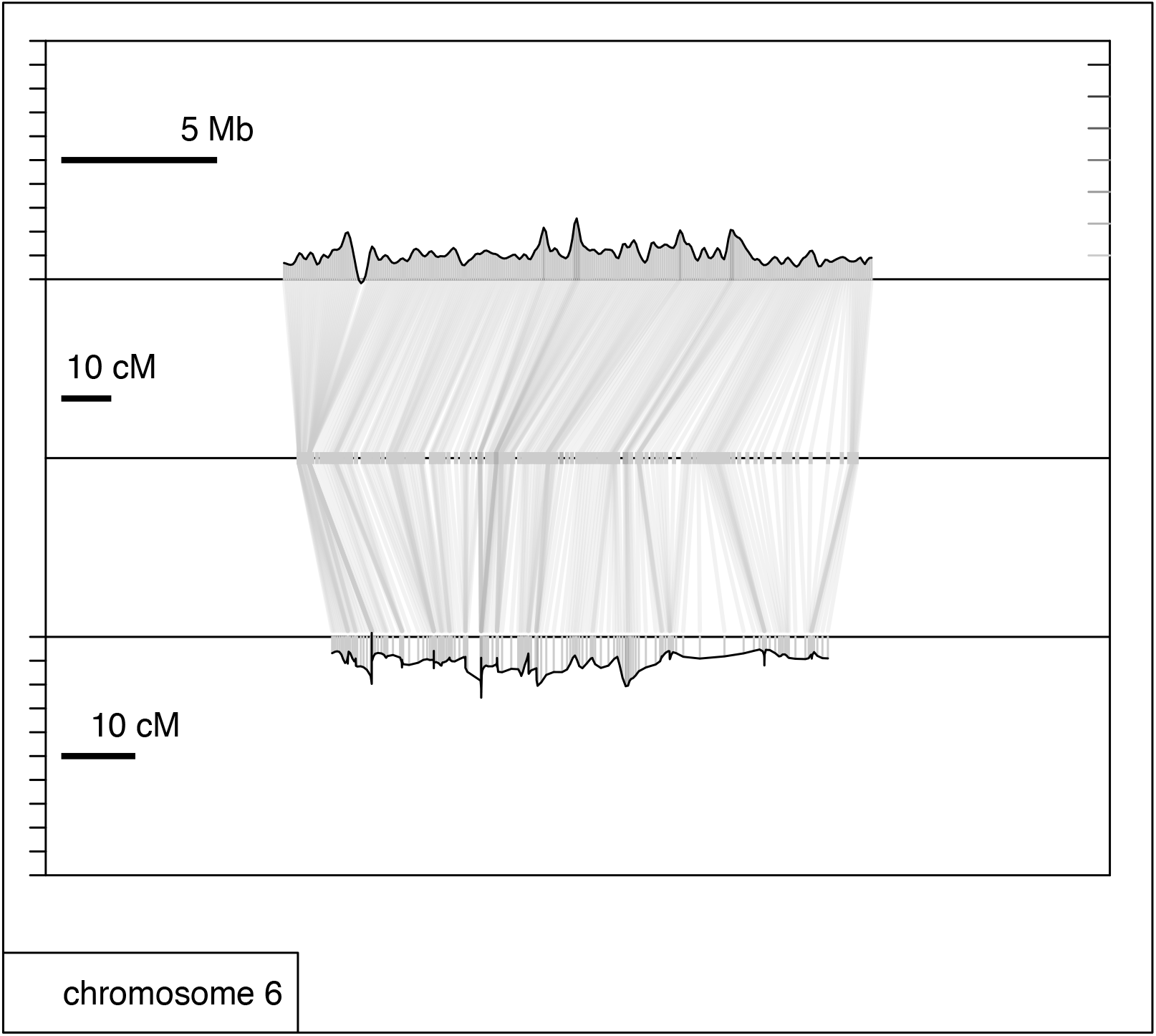
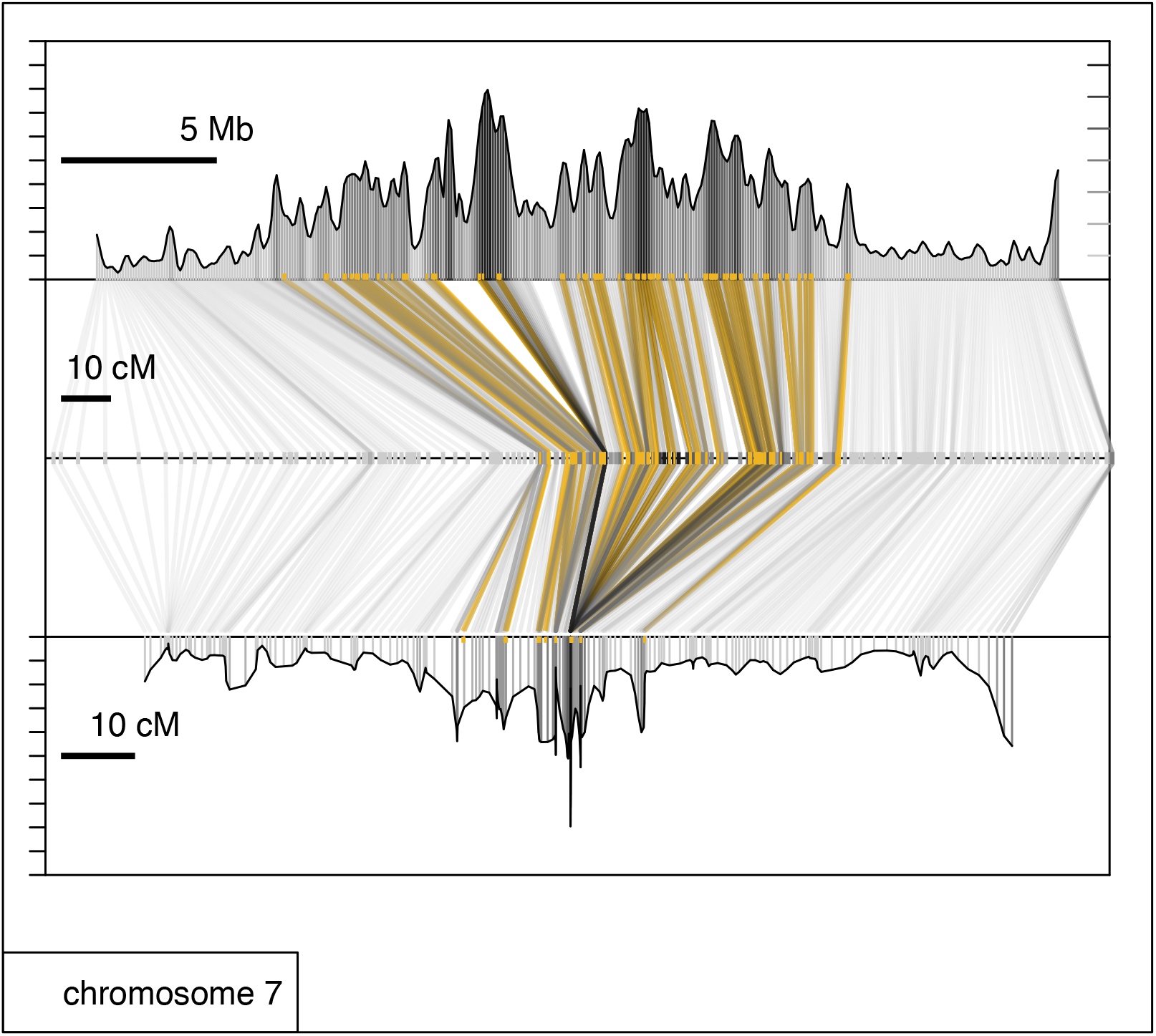
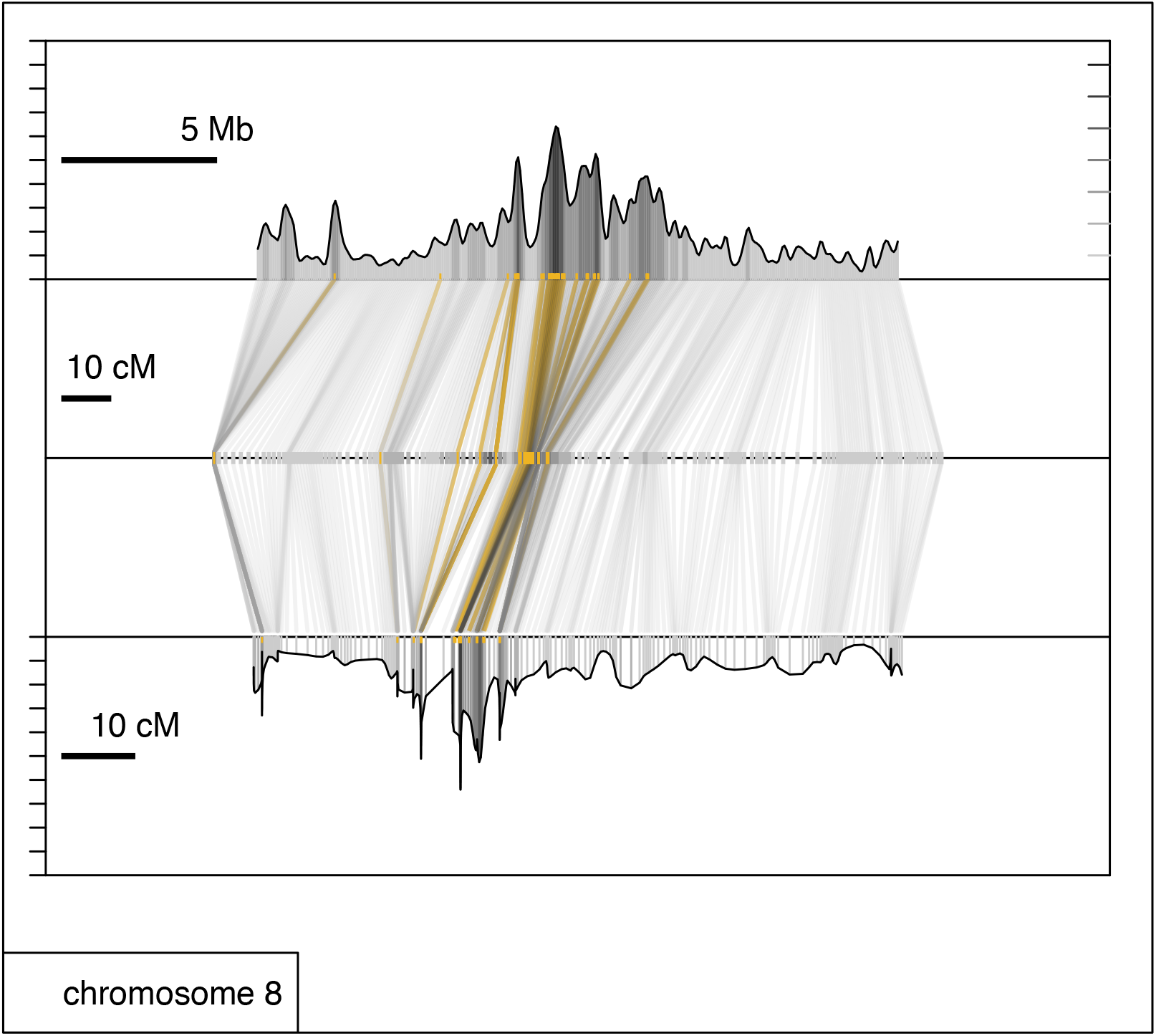
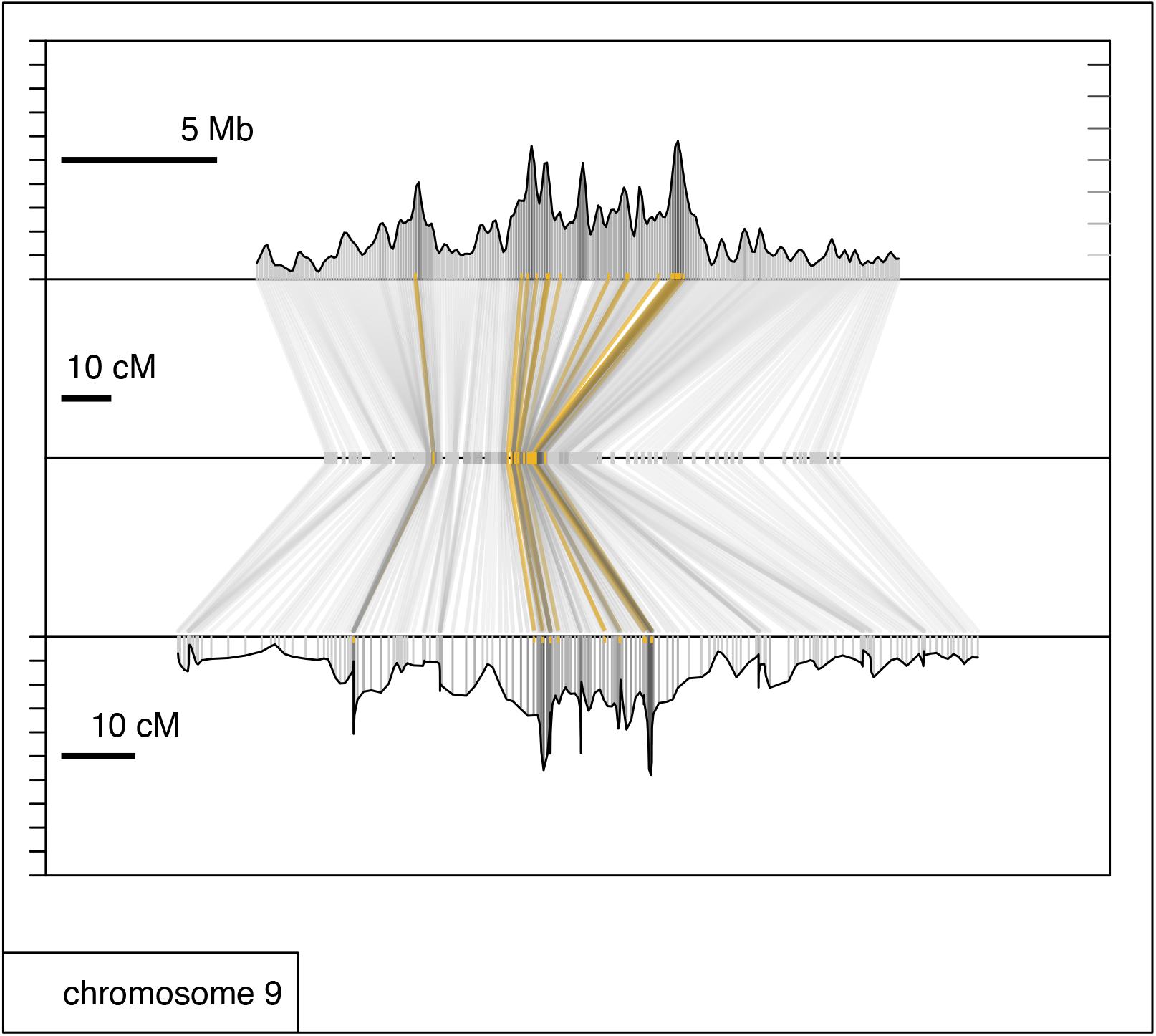
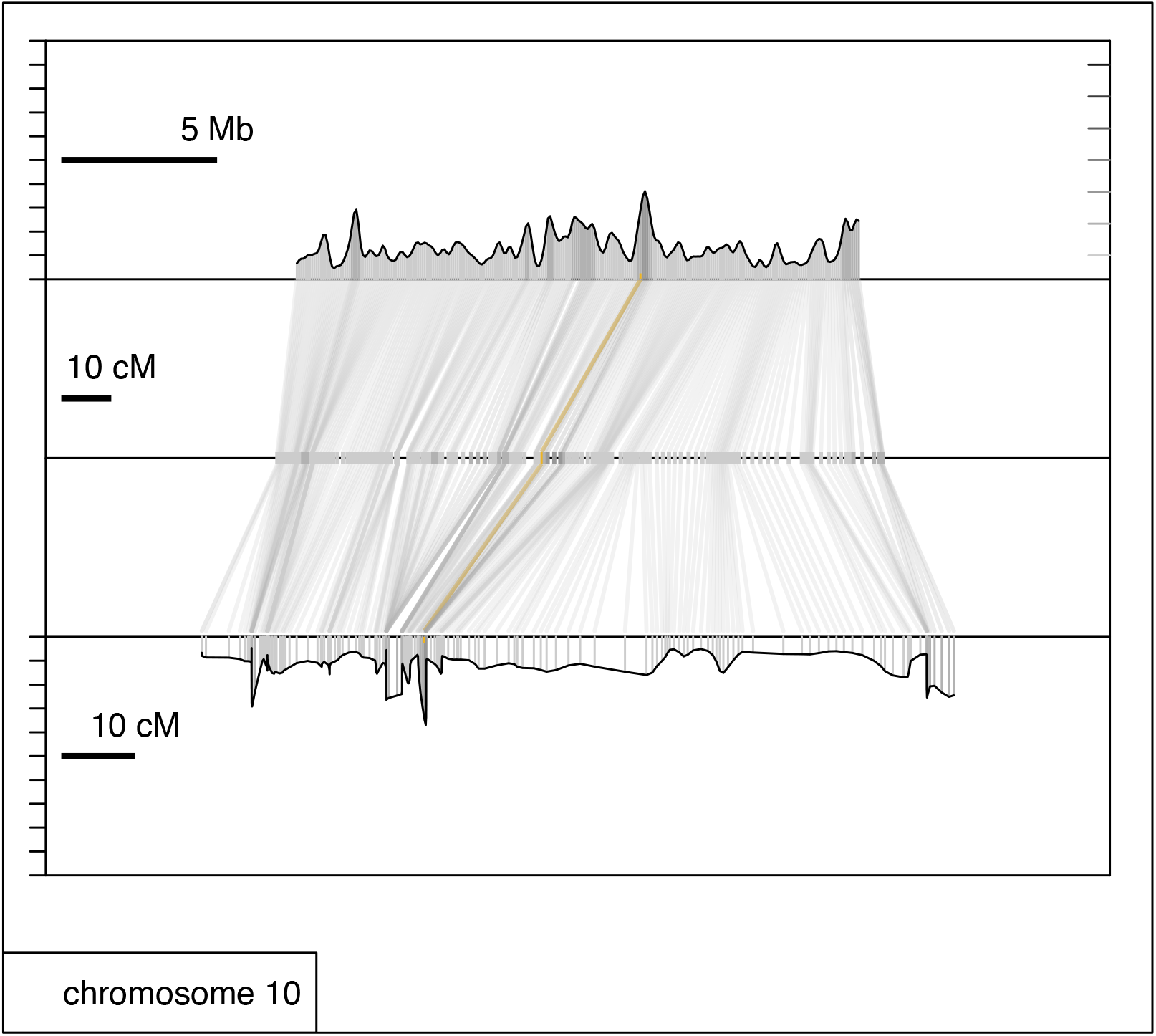
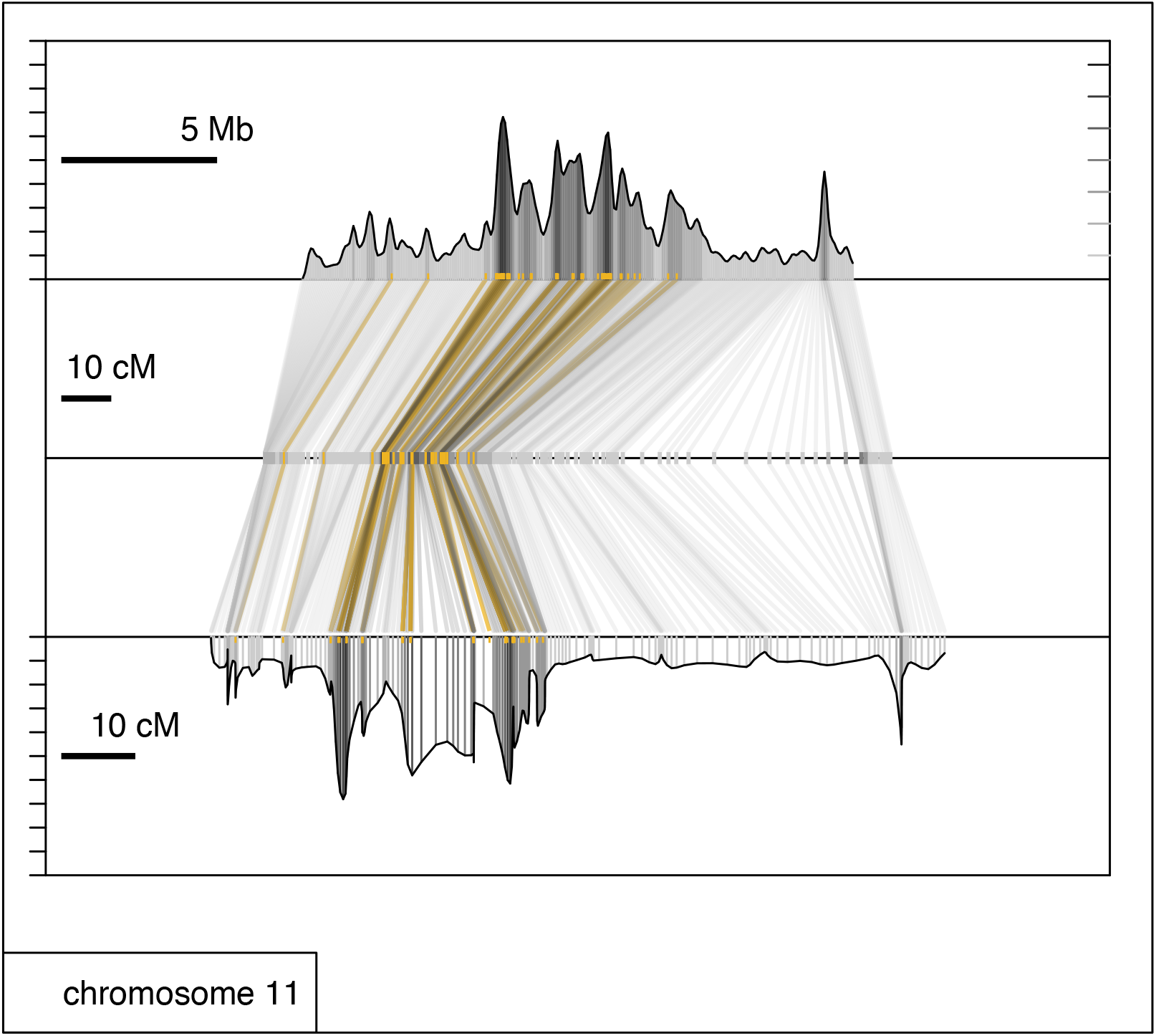
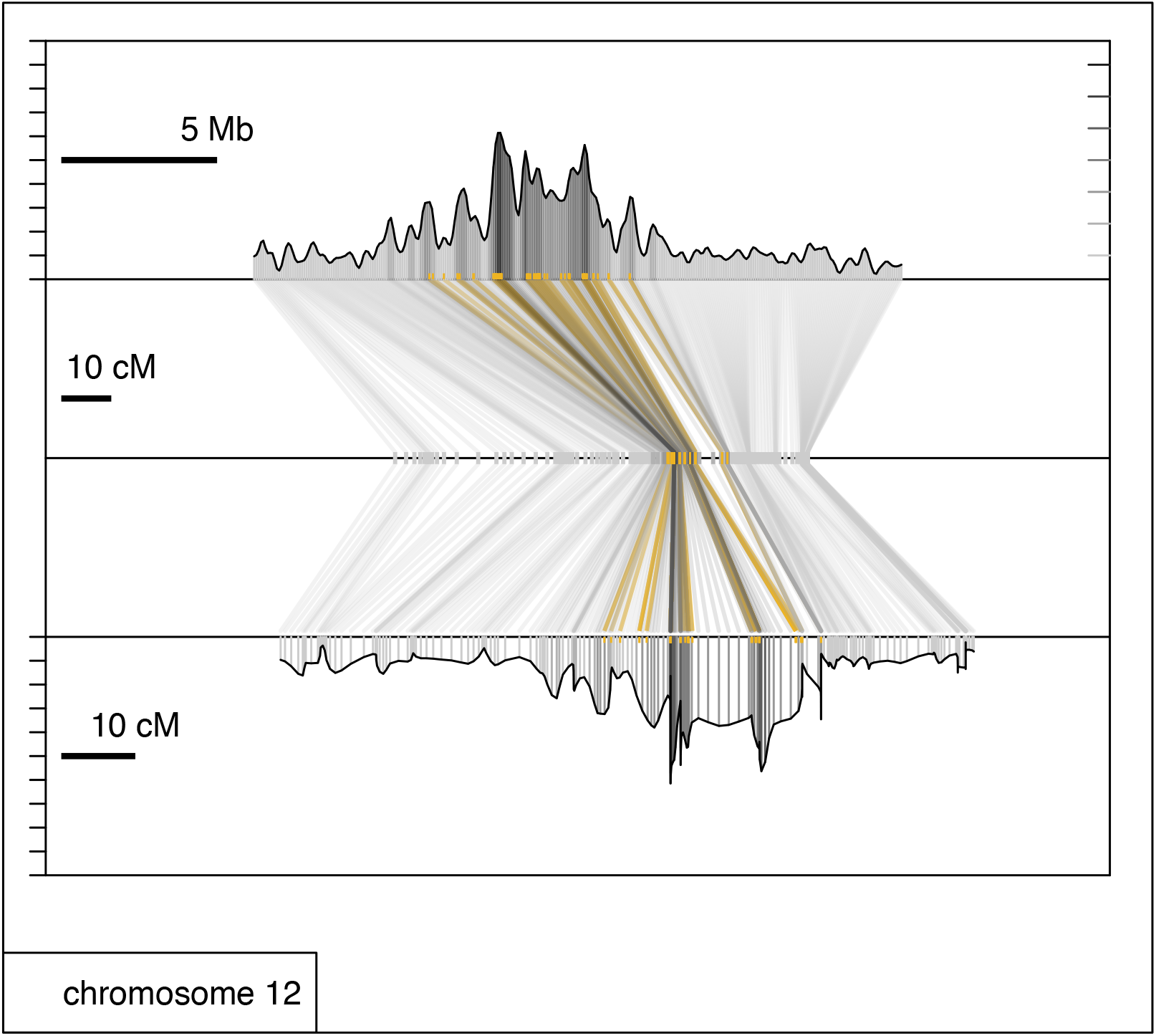
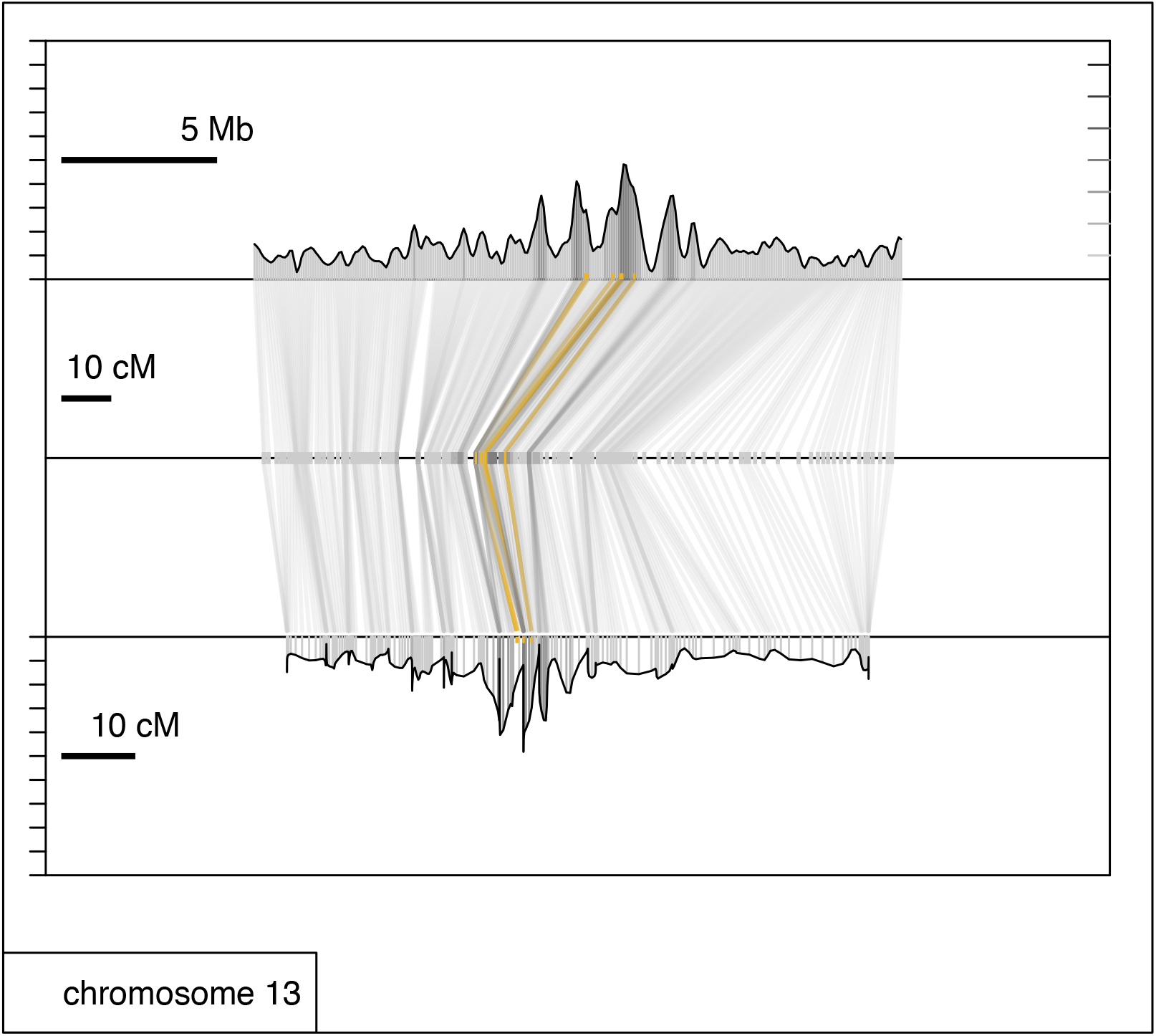
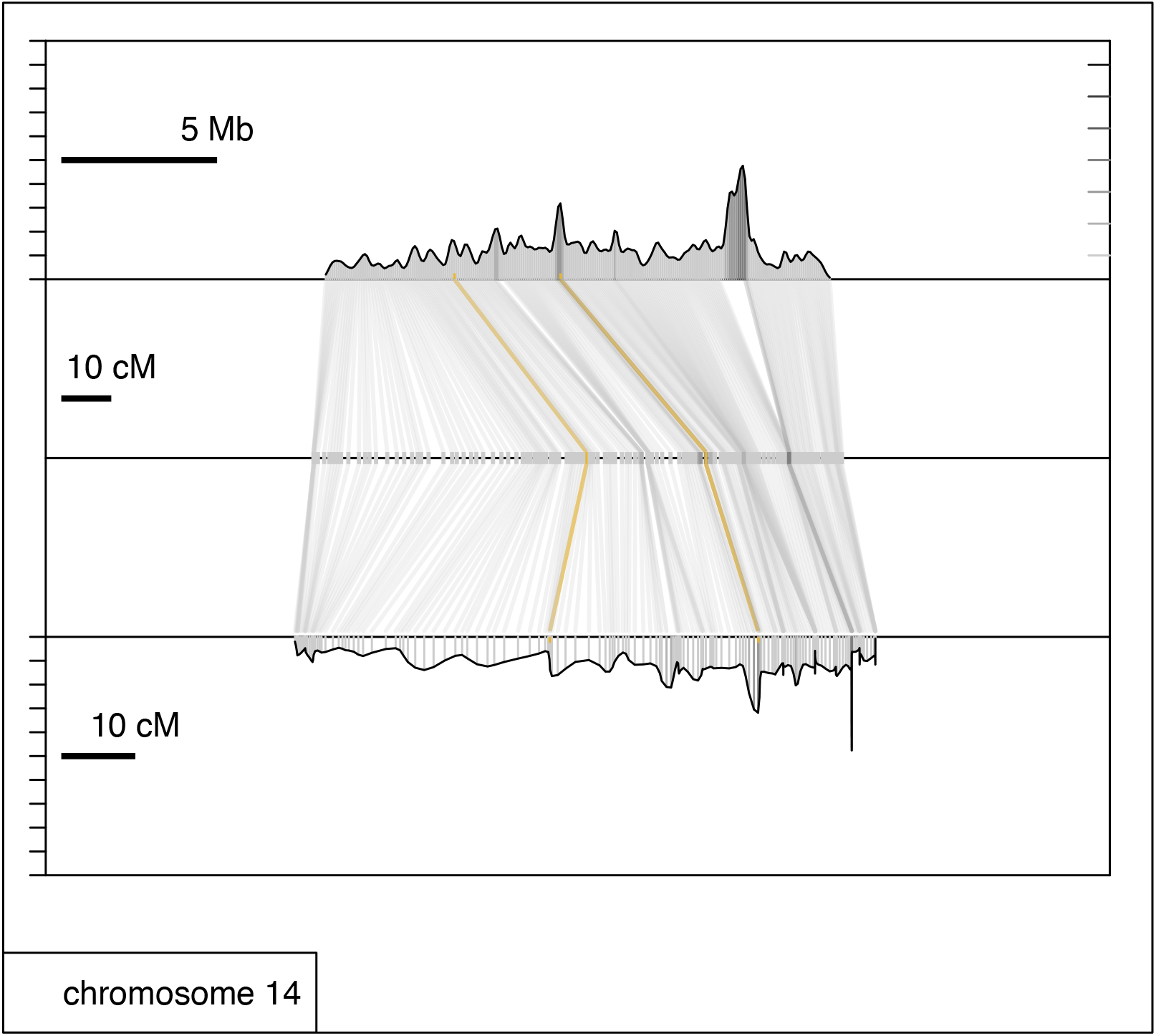
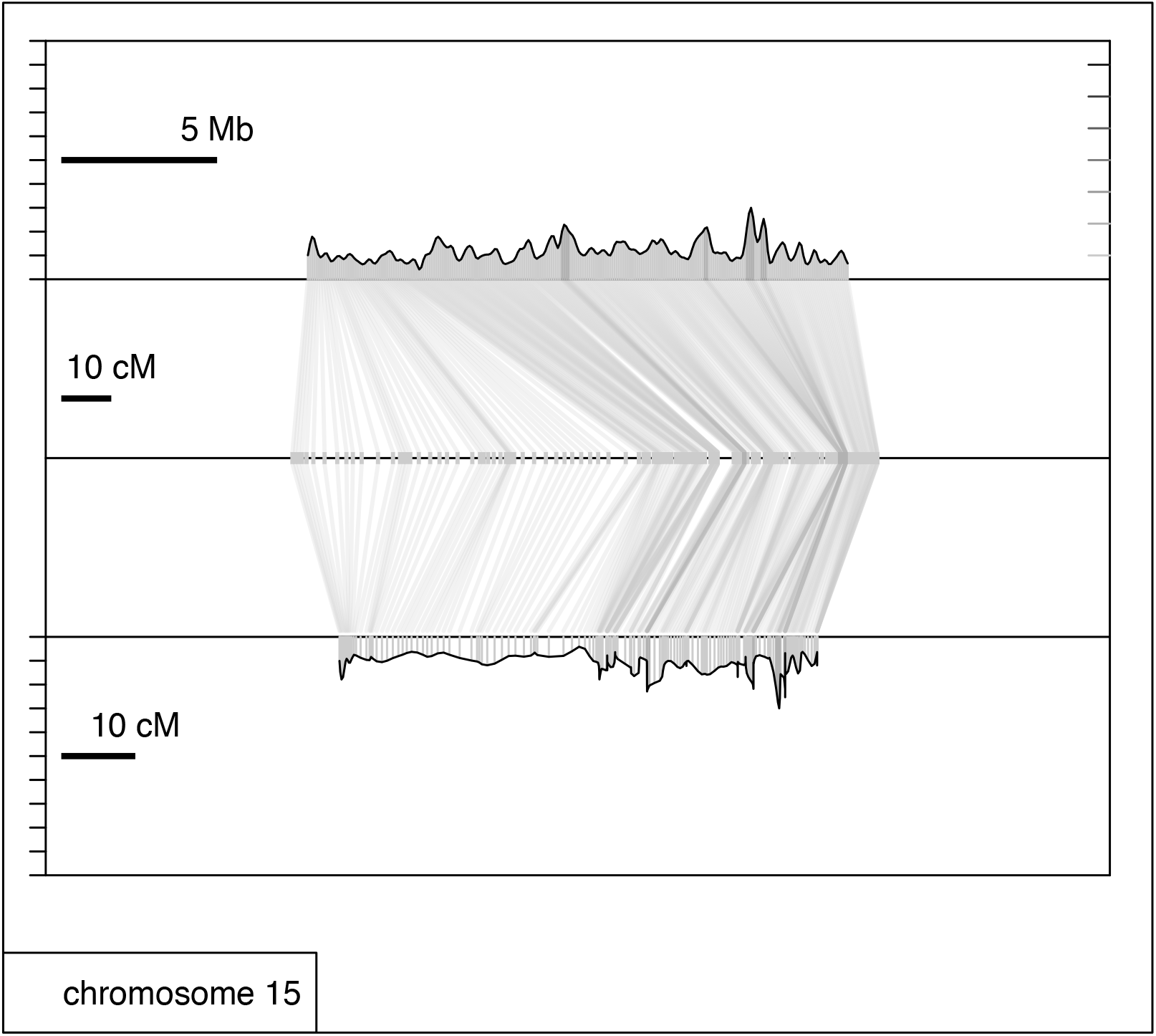
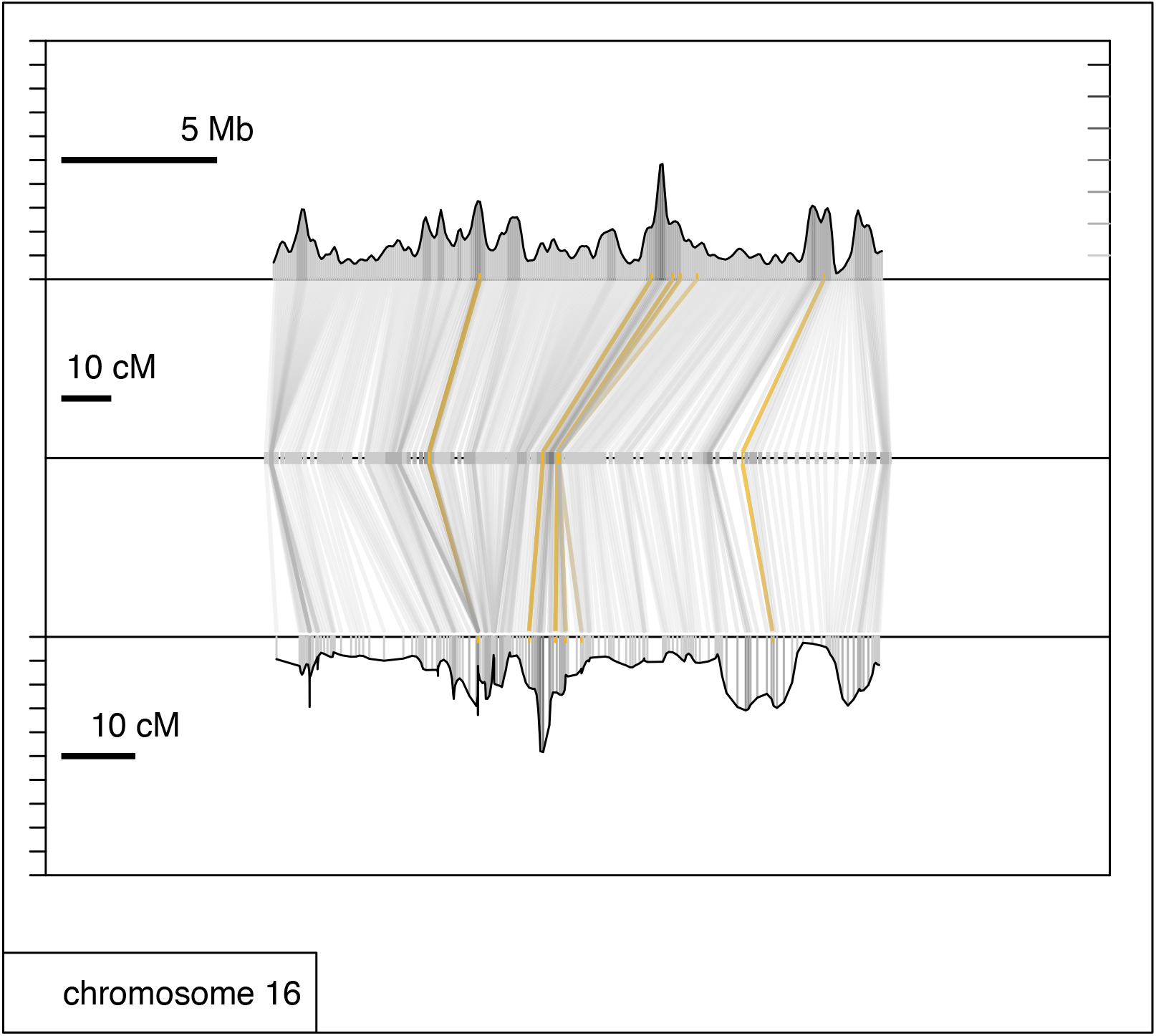
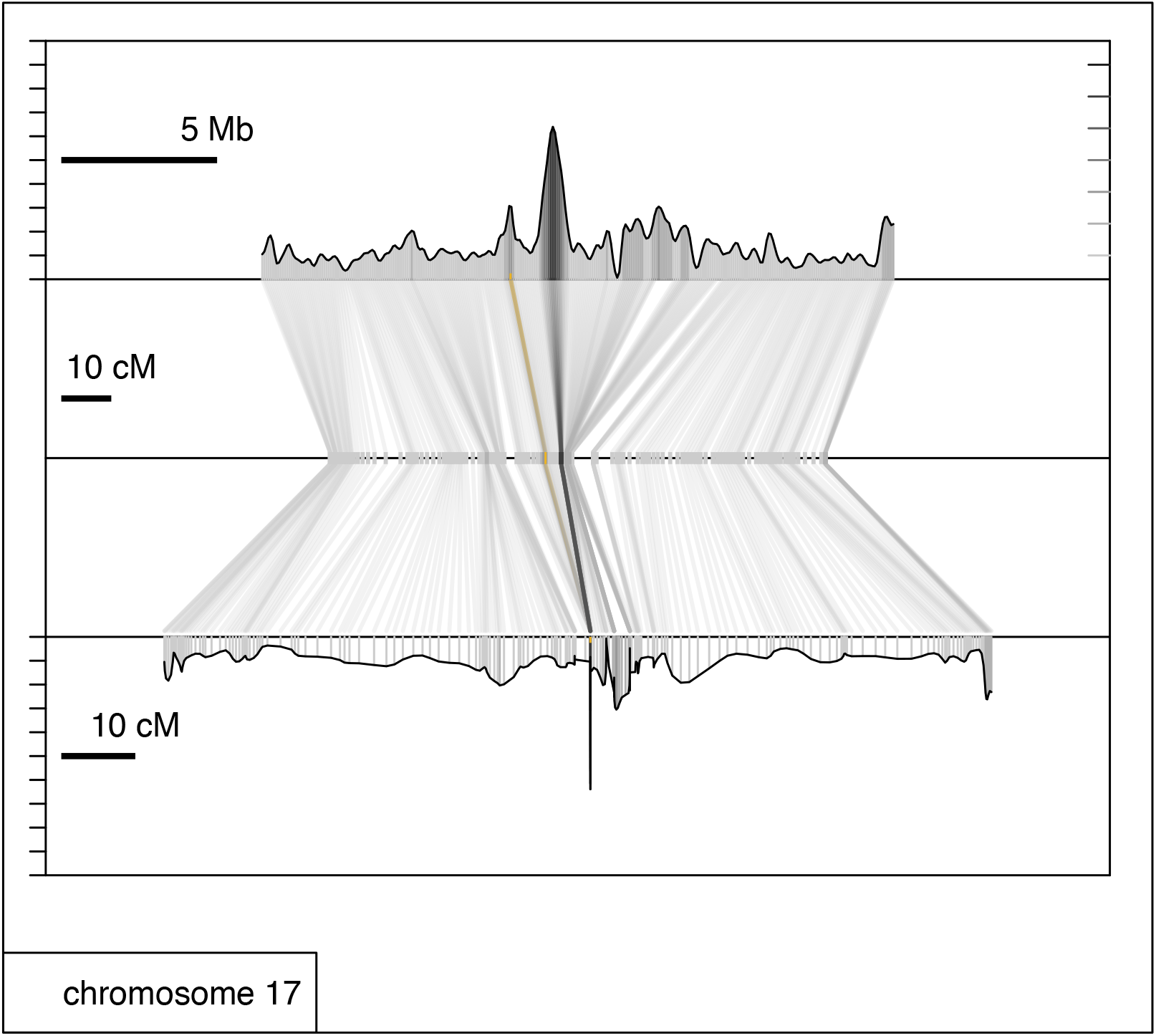
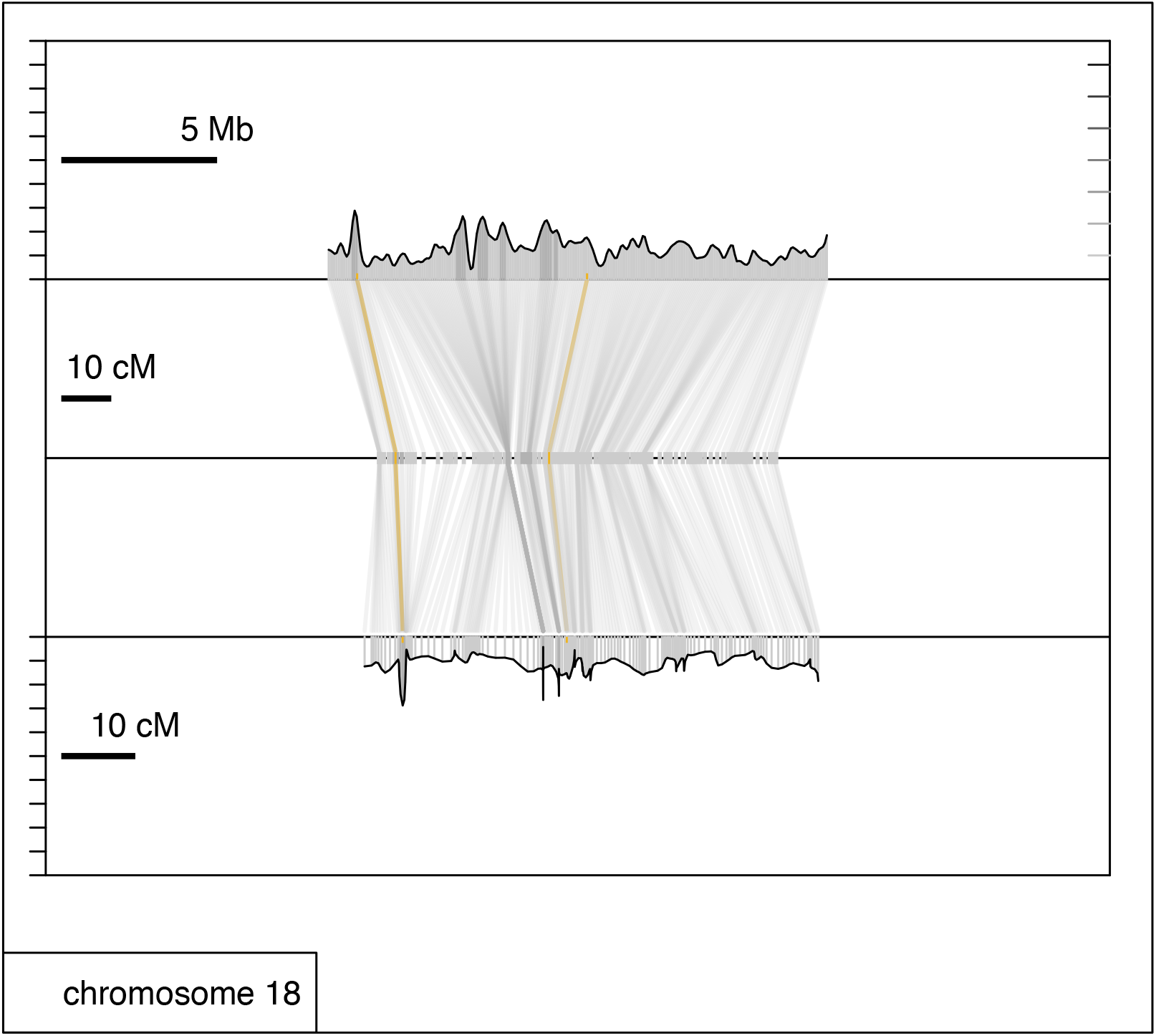
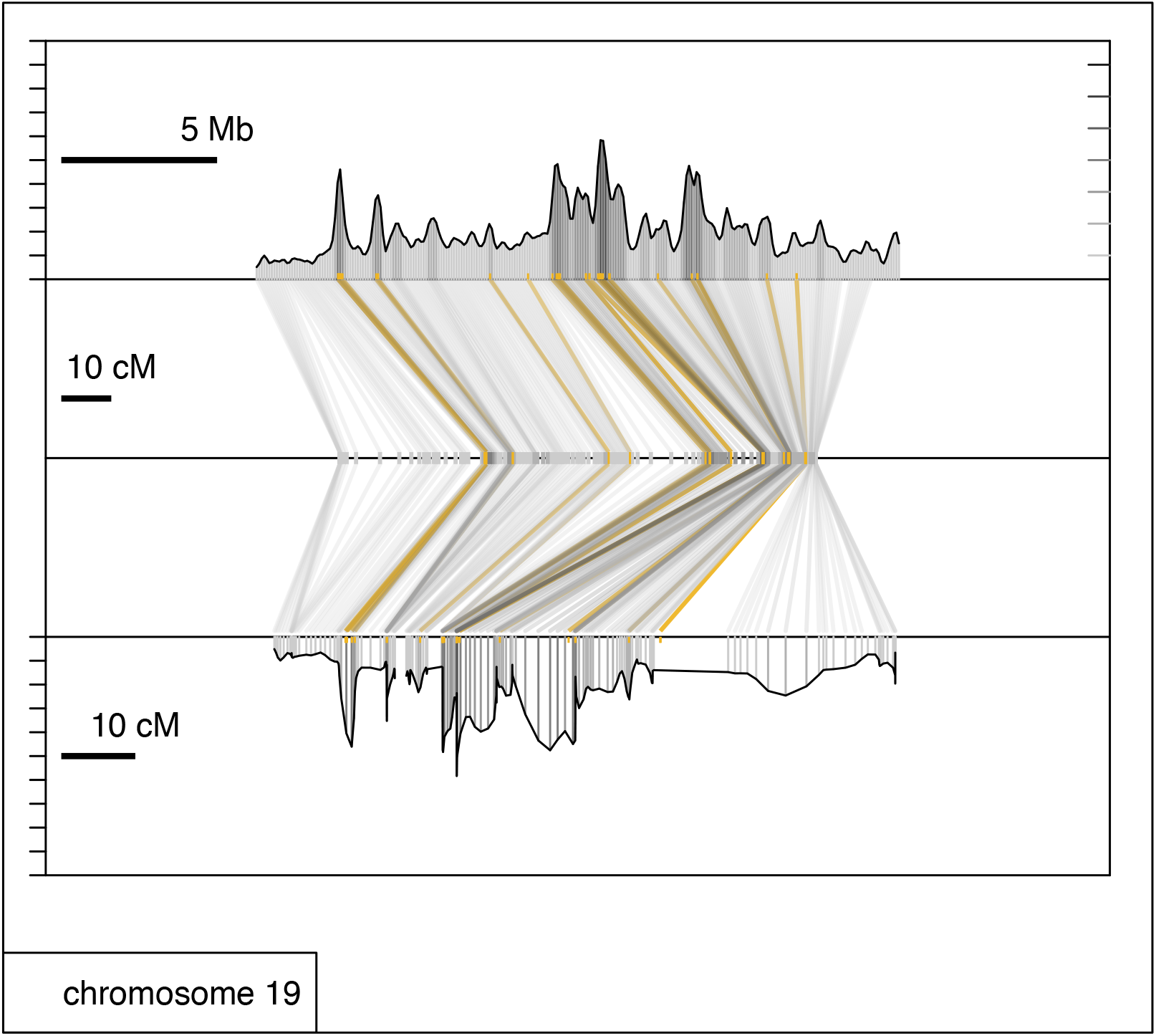
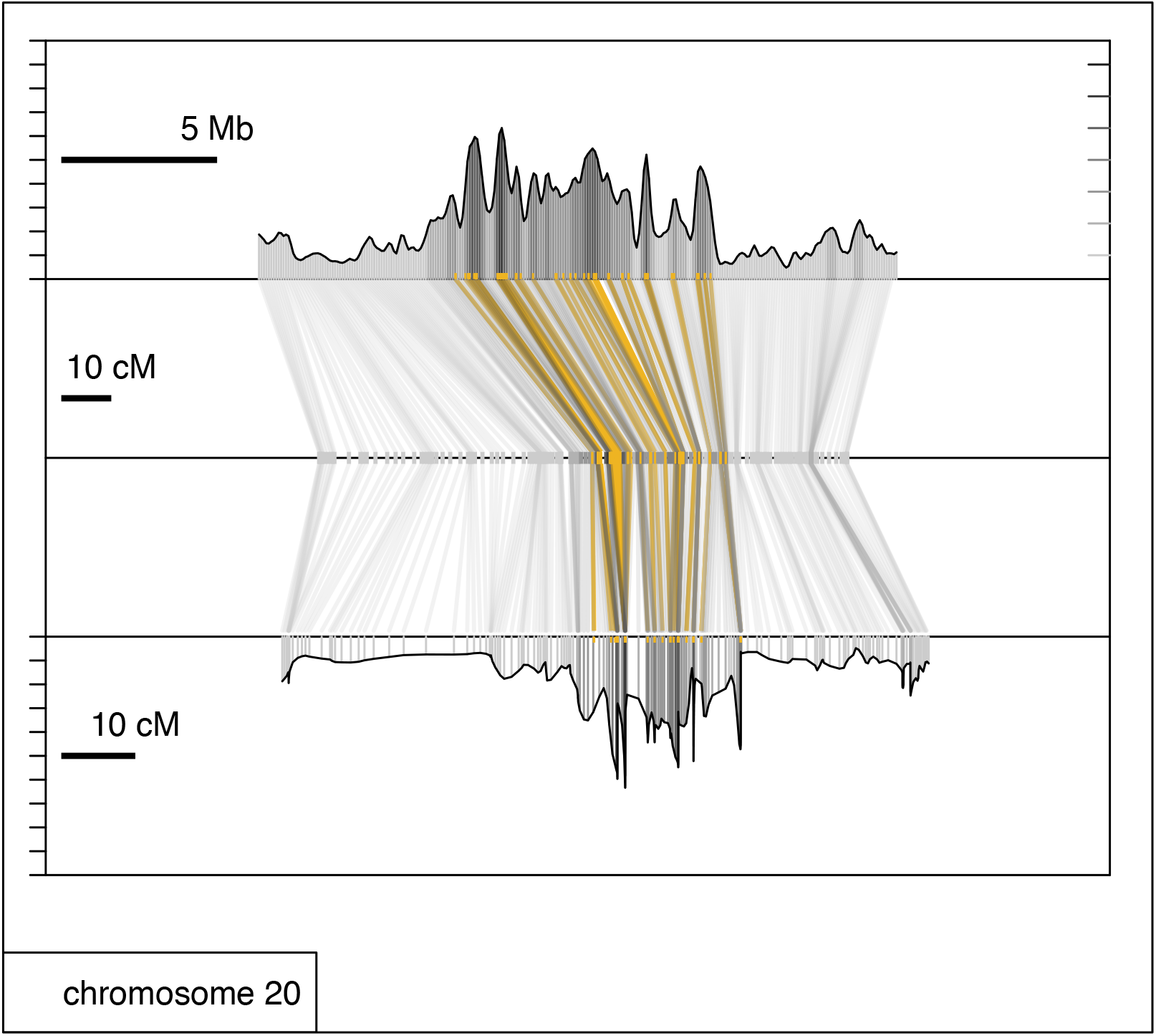
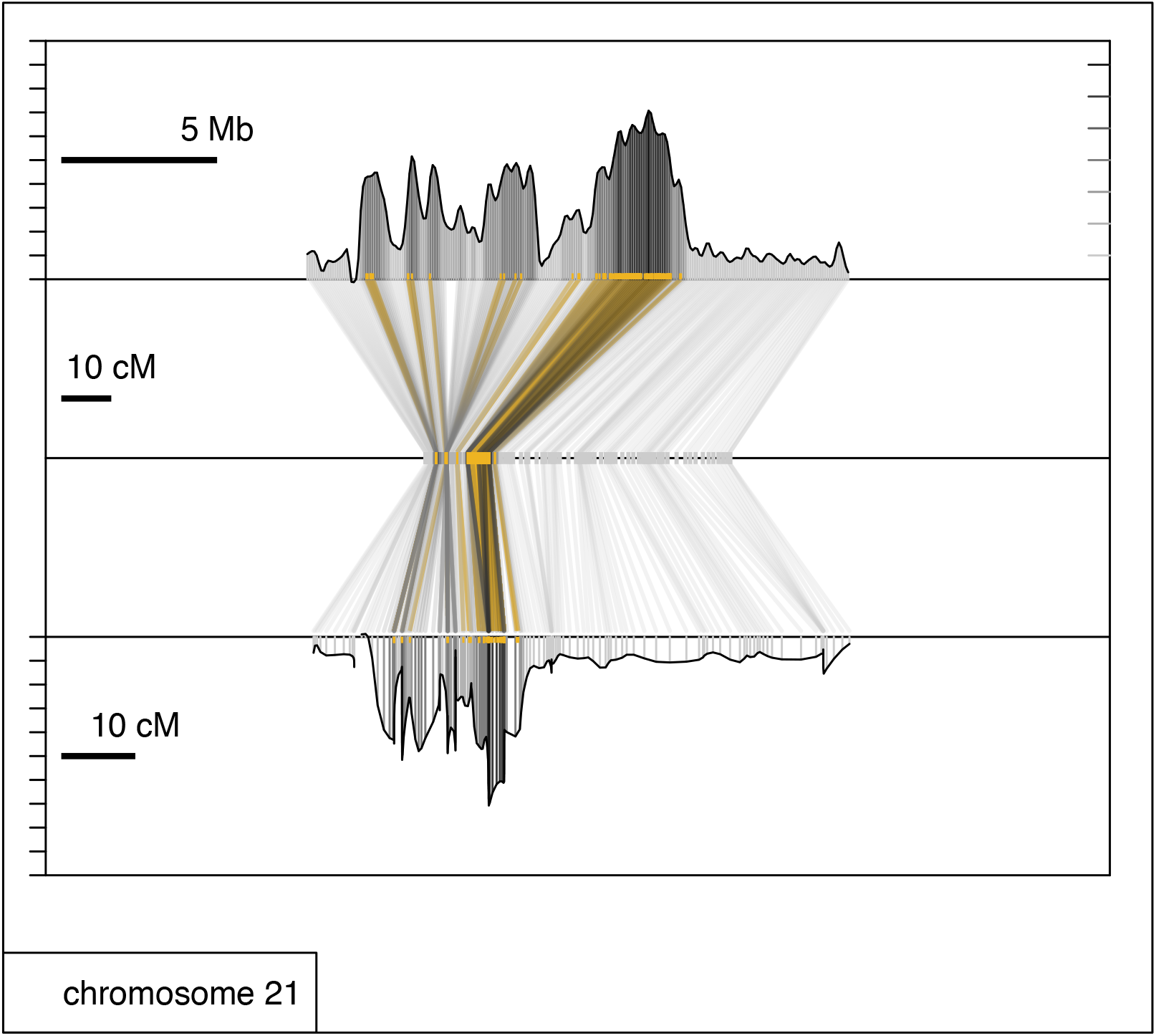

## Literature Cited

Aeschbacher, S., J. P. Selby, J. H. Willis and G. Coop, 2017 Population-genomic inference of the strength and timing of selection against gene flow. Proc Natl Acad Sci U S A 114: 7061–7066.

Angert, A. L., M. Bayly, S. N. Sheth and J. R. Paul, 2018 Testing range-limit hypotheses using range-wide habitat suitability and occupancy for the scarlet monkeyflower (Erythranthe cardinalis). Am Nat 191: E76-E89.

Aronesty, E., 2011 ea-utils: Command-line tools for processing biological sequencing data.

Baird, N. A., P. D. Etter, T. S. Atwood, M. C. Currey, A. L. Shiver et al., 2008 Rapid SNP discovery and genetic mapping using sequenced RAD markers. PLoS One 3: e3376.

Barrett, R. D., S. M. Rogers and D. Schluter, 2008 Natural selection on a major armor gene in threespine stickleback. Science 322: 255–257.

Bassham, S., J. Catchen, E. Lescak, F. A. von Hippel and W. A. Cresko, 2018 Repeated selection of alternatively adapted haplotypes creates sweeping genomic remodeling in stickleback. Genetics 209: 921–939.

Begun, D. J., and C. F. Aquadro, 1992 Levels of naturally occurring DNA polymorphism correlate with recombination rates in D. melanogaster. Nature 356: 519–520.

Begun, D. J., A. K. Holloway, K. Stevens, L. W. Hillier, Y. P. Poh et al., 2007 Population genomics: whole-genome analysis of polymorphism and divergence in Drosophila simulans. PLoS Biol 5: e310.

Bell, M. A., and S. A. Foster, 1994 Introduction to the evolutionary biology of the threespine stickleback, pp. 1-27 in The Evolutionary Biology of the Threespine Stickleback, edited by M. A. Bell and S. A. Foster. Oxford University Press, New York.

Bergland, A. O., E. L. Behrman, K. R. O’Brien, P. S. Schmidt and D. A. Petrov, 2014 Genomic Evidence of Rapid and Stable Adaptive Oscillations over Seasonal Time Scales in Drosophila. PLoS Genet 10.

Berner, D., A. C. Grandchamp and A. P. Hendry, 2009 Variable progress toward ecological speciation in parapatry: stickleback across eight lake-stream transitions. Evolution 63: 1740–1753.

Burri, R., A. Nater, T. Kawakami, C. F. Mugal, P. I. Olason et al., 2015 Linked selection and recombination rate variation drive the evolution of the genomic landscape of differentiation across the speciation continuum of Ficedula flycatchers. Genome Res 25: 1656–1665.

Caldera, E. J., and D. I. Bolnick, 2008 Effects of colonization history and landscape structure on genetic variation within and among threespine stickleback (Gasterosteus aculeatus) populations in a single watershed. Evol Ecol Res 10: 575–598.

Carneiro, M., N. Ferrand and M. W. Nachman, 2009 Recombination and speciation: loci near centromeres are more differentiated than loci near telomeres between subspecies of the European rabbit (Oryctolagus cuniculus). Genetics 181: 593–606.

Catchen, J., S. Bassham, T. Wilson, M. Currey, C. O’Brien et al., 2013a The population structure and recent colonization history of Oregon threespine stickleback determined using restriction-site associated DNA-sequencing. Mol Ecol 22: 2864–2883.

Catchen, J., P. A. Hohenlohe, S. Bassham, A. Amores and W. A. Cresko, 2013b Stacks: an analysis tool set for population genomics. Mol Ecol 22: 3124–3140.

Catchen, J. M., A. Amores, P. Hohenlohe, W. Cresko and J. H. Postlethwait, 2011 Stacks: building and genotyping loci de novo from short-read sequences. G3-Genes Genomes Genetics 1: 171–182.

Charlesworth, B., D. Charlesworth and N. H. Barton, 2003 The effects of genetic and geographic structure on neutral variation. Ann Rev Ecol Evol Systemat 34: 99–125.

Charlesworth, B., M. T. Morgan and D. Charlesworth, 1993 The effect of deleterious mutations on neutral molecular variation. Genetics 134: 1289–1303.

Charlesworth, B., M. Nordborg and D. Charlesworth, 1997 The effects of local selection, balanced polymorphism and background selection on equilibrium patterns of genetic diversity in subdivided populations. Genet Res 70: 155–174.

Clausen, J., W. M. Keck and W. M. Hiesey, 1941 Regional differentiation in plant species. Am Nat 75: 231–250.

Colosimo, P. F., K. E. Hosemann, S. Balabhadra, G. Villarreal, Jr., M. Dickson et al., 2005 Widespread parallel evolution in sticklebacks by repeated fixation of Ectodysplasin alleles. Science 307: 1928–1933.

Cresko, W. A., A. Amores, C. Wilson, J. Murphy, M. Currey et al., 2004 Parallel genetic basis for repeated evolution of armor loss in Alaskan threespine stickleback populations. Proc Natl Acad Sci U S A 101: 6050–6055.

Cruickshank, T. E., and M. W. Hahn, 2014 Reanalysis suggests that genomic islands of speciation are due to reduced diversity, not reduced gene flow. Mol Ecol 23: 3133–3157.

Cutter, A. D., and B. A. Payseur, 2013 Genomic signatures of selection at linked sites: unifying the disparity among species. Nat Rev Genet 14: 262–274.

Deagle, B. E., F. C. Jones, Y. F. Chan, D. M. Absher, D. M. Kingsley et al., 2012 Population genomics of parallel phenotypic evolution in stickleback across stream-lake ecological transitions. Proc R Soc B-Biol Sci 279: 1277–1286.

Defaveri, J., T. Shikano, Y. Shimada and J. Merilä, 2013 High degree of genetic differentiation in marine three-spined sticklebacks (Gasterosteus aculeatus). Mol Ecol 22: 4811–4828.

Dobzhansky, T., and M. L. Queal, 1938 Genetics of Natural Populations. II. Genic Variation in Populations of Drosophila Pseudoobscura Inhabiting Isolated Mountain Ranges. Genetics 23: 463–484.

Drummond, A. J., and A. Rambaut, 2007 BEAST: Bayesian evolutionary analysis by sampling trees. BMC Evol Biol 7: 214.

Drummond, A. J., M. A. Suchard, D. Xie and A. Rambaut, 2012 Bayesian phylogenetics with BEAUti and the BEAST 1.7. Mol Biol Evol 29: 1969–1973.

Ellegren, H., L. Smeds, R. Burri, P. I. olason, N. Backström et al., 2012 The genomic landscape of species divergence in Ficedula flycatchers. Nature: 1–5.

Elyashiv, E., S. Sattath, T. T. Hu, A. Strutsovsky, G. McVicker et al., 2016 A genomic map of the effects of linked selection in Drosophila. PLoS Genet 12: e1006130.

Endler, J. A., 1977 Geographic variation, speciation, and clines, pp. Princeton University Press, Princeton, NJ.

Fay, J. C., and C. I. Wu, 2000 Hitchhiking under positive Darwinian selection. Genetics 155: 1405–1413.

Fontaine, M. C., J. B. Pease, A. Steele, R. M. Waterhouse, D. E. Neafsey et al., 2015 Mosquito genomics. Extensive introgression in a malaria vector species complex revealed by phylogenomics. Science 347: 1258524.

Geraldes, A., P. Basset, K. L. Smith and M. W. Nachman, 2011 Higher differentiation among subspecies of the house mouse (Mus musculus) in genomic regions with low recombination. Mol Ecol 20: 4722–4736.

Gillespie, J. H., 2000 Genetic drift in an infinite population: The pseudohitchhiking model. Genetics 155: 909–919.

Glazer, A. M., E. E. Killingbeck, T. Mitros, D. S. Rokhsar and C. T. Miller, 2015 Genome Assembly Improvement and Mapping Convergently Evolved Skeletal Traits in Sticklebacks with Genotyping-by-Sequencing. G3-Genes Genomes Genetics 5: 1463–1472.

Hahn, M. W., 2008 Toward a selection theory of molecular evolution. Evolution 62: 255–265.

Haller, B. C., and P. W. Messer, 2017 SLiM 2: flexible, interactive forward genetic simulations. Mol Biol Evol 34: 230–240.

Hermisson, J., and P. S. Pennings, 2005 Soft sweeps: Molecular population genetics of adaptation from standing genetic variation. Genetics 169: 2335–2352.

Hoekstra, H. E., K. E. Drumm and M. W. Nachman, 2004 Ecological genetics of adaptive color polymorphism in pocket mice: geographic variation in selected and neutral genes. Evolution 58: 1329–1341.

Hohenlohe, P. A., S. Bassham, M. Currey and W. A. Cresko, 2012 Extensive linkage disequilibrium and parallel adaptive divergence across threespine stickleback genomes. Philos Trans R Soc Lond B Biol Sci 367: 395–408.

Hohenlohe, P. A., S. Bassham, P. D. Etter, N. Stiffler, E. A. Johnson et al., 2010 Population genomics of parallel adaptation in threespine stickleback using sequenced RAD tags. PLoS Genet 6: e1000862.

Hubby, J. L., and R. C. Lewontin, 1966 A molecular approach to the study of genic heterozygosity in natural populations. I. The number of alleles at different loci in Drosophila pseudoobscura. Genetics 54: 577–594.

Hudson, R. R., M. Slatkin and W. P. Maddison, 1992 Estimation of levels of gene flow from DNA-sequence data. Genetics 132: 583–589.

Jones, F. C., M. G. Grabherr, Y. F. Chan, P. Russell, E. Mauceli et al., 2012 The genomic basis of adaptive evolution in threespine sticklebacks. Nature 484: 55–61.

Jones, M. R., L. S. Mills, P. C. Alves, C. M. Callahan, J. M. Alves et al., 2018 Adaptive introgression underlies polymorphic seasonal camouflage in snowshoe hares. Science 360: 1355–1358.

Kaeuffer, R., C. L. Peichel, D. I. Bolnick and A. P. Hendry, 2012 Parallel and nonparallel aspects of ecological, phenotypic, and genetic divergence across replicate population pairs of lake and stream stickleback. Evolution 66: 402–418.

Kern, A. D., and M. W. Hahn, 2018 The neutral theory in light of natural selection. Mol Biol Evol 35: 1366–1371.

Kirkpatrick, M., and N. Barton, 2006 Chromosome inversions, local adaptation and speciation. Genetics 173: 419–434.

Kitano, J., D. I. Bolnick, D. A. Beauchamp, M. M. Mazur, S. Mori et al., 2008 Reverse evolution of armor plates in the threespine stickleback. Curr Biol 18: 769–774.

Langley, C. H., K. Stevens, C. Cardeno, Y. C. Lee, D. R. Schrider et al., 2012 Genomic variation in natural populations of Drosophila melanogaster. Genetics 192: 533–598.

Leaver, S. D., and T. E. Reimchen, 2012 Abrupt changes in defence and trophic morphology of the giant threespine stickleback (Gasterosteus sp.) following colonization of a vacant habitat. Biol J Linn Soc 107: 494–509.

Lenormand, T., 2002 Gene flow and the limits to natural selection. Trends in Ecology & Evolution 17: 183–189.

Lenormand, T., D. Bourguet, T. Guillemaud and M. Raymond, 1999 Tracking the evolution of insecticide resistance in the mosquito Culex pipiens. Nature 400: 861–864.

Lescak, E. A., S. L. Bassham, J. Catchen, O. Gelmond, M. L. Sherbick et al., 2015 Evolution of stickleback in 50 years on earthquake-uplifted islands. Proc Natl Acad Sci U S A 112: E7204–7212.

Lewontin, R. C., and J. L. Hubby, 1966 A molecular approach to study of genic heterozygosity in natural populations. II. Amount of variation and degree of heterozygosity in natural populations of Drosophila pseudoobscura. Genetics 54: 595–609.

Li, L., M. Jean and F. Belzile, 2006 The impact of sequence divergence and DNA mismatch repair on homeologous recombination in Arabidopsis. Plant J 45: 908–916.

Marques, D. A., K. Lucek, J. I. Meier, S. Mwaiko, C. E. Wagner et al., 2016 Genomics of rapid incipient speciation in sympatric threespine stickleback. PLoS Genet 12: e1005887.

Maynard Smith, J., and J. Haigh, 1974 The hitch-hiking effect of a favourable gene. Genetical Research 23: 23–35.

McKinnon, J. S., and H. D. Rundle, 2002 Speciation in nature: the threespine stickleback model systems. TREE 17: 480–488.

Meier, J. I., D. A. Marques, S. Mwaiko, C. E. Wagner, L. Excoffier et al., 2017 Ancient hybridization fuels rapid cichlid fish adaptive radiations. Nat Comm 8: 14363.

Mettler, L. E., R. A. Voelker and T. Mukai, 1977 Inversion clines in populations of Drosophila melanogaster. Genetics 87: 169–176.

Modrich, P., and R. Lahue, 1996 Mismatch repair in replication fidelity, genetic recombination, and cancer biology. Ann Rev Biochem 65: 101–133.

Nachman, M. W., 2002 Variation in recombination rate across the genome: evidence and implications. Curr Opinion Genet Devel 12: 657–663.

Nei, M., 1987 Molecular evolutionary genetics. Columbia university press.

Nelson, T. C., and W. A. Cresko, 2018 Ancient genomic variation underlies repeated ecological adaptation in young stickleback populations. Evol Lett 2: 9–21.

Nielsen, R., S. Williamson, Y. Kim, M. J. Hubisz, A. G. Clark et al., 2005 Genomic scans for selective sweeps using SNP data. Genome Res 15: 1566–1575.

Nosil, P., and B. J. Crespi, 2006 Experimental evidence that predation promotes divergence in adaptive radiation. Proc Natl Acad Sci U S A 103: 9090–9095.

Opperman, R., E. Emmanuel and A. A. Levy, 2004 The effect of sequence divergence on recombination between direct repeats in Arabidopsis. Genetics 168: 2207.

Paradis, E., J. Claude and K. Strimmer, 2004 APE: Analyses of phylogenetics and evolution in R language. Bioinformat 20: 289–290.

Pardo-Diaz, C., C. Salazar, S. W. Baxter, C. Merot, W. Figueiredo-Ready et al., 2012 Adaptive introgression across species boundaries in Heliconius butterflies. PLoS Genet 8.

Pfeifer, B., U. Wittelsburger, S. E. Ramos-Onsins and M. J. Lercher, 2014 PopGenome: an efficient Swiss army knife for population genomic analyses in R. Mol Biol Evol 31: 1929–1936.

Popescu, A. A., K. T. Huber and E. Paradis, 2012 ape 3.0: New tools for distance-based phylogenetics and evolutionary analysis in R. Bioinformat 28: 1536–1537.

Rastas, P., F. C. Calboli, B. Guo, T. Shikano and J. Merila, 2015 Construction of ultradense linkage maps with Lep-MAP2: stickleback F2 recombinant crosses as an example. Genome Biol Evol 8: 78–93.

Reimchen, T. E., 1994 Predators and morphological evolution in threespine stickleback, pp. 240–276 in The Evolutionary Biology of the Threespine Stickleback, edited by M. A. Bell and S. A. Foster. Oxford University Press, New York.

Reimchen, T. E., C. Bergstrom and P. Nosil, 2013 Natural selection and the adaptive radiation of Haida Gwaii stickleback. Evol Ecol Res 15: 241–269.

Roesti, M., S. Gavrilets, A. P. Hendry, W. Salzburger and D. Berner, 2014 The genomic signature of parallel adaptation from shared genetic variation. Mol Ecol 23: 3944–3956.

Roesti, M., B. Kueng, D. Moser and D. Berner, 2015 The genomics of ecological vicariance in threespine stickleback fish. Nat Comm 6: 8767.

Roesti, M., D. Moser and D. Berner, 2013 Recombination in the threespine stickleback genome - Patterns and consequences. Mol Ecol 22: 3014–3027.

Samuk, K., G. L. Owens, K. E. Delmore, S. E. Miller, D. J. Rennison et al., 2017 Gene flow and selection interact to promote adaptive divergence in regions of low recombination. Mol Ecol 26: 4378–4390.

Sardell, J. M., C. D. Cheng, A. J. Dagilis, A. Ishikawa, J. Kitano et al., 2018 Sex Differences in Recombination in Sticklebacks. G3-Genes Genomes Genetics 8: 1971–1983.

Scheet, P., and M. Stephens, 2006 A fast and flexible statistical model for large-scale population genotype data: applications to inferring missing genotypes and haplotypic phase. Am J Hum Genet 78: 629–644.

Schluter, D., and G. L. Conte, 2009 Genetics and ecological speciation. Proc Natl Acad Sci U S A 106: 9955–9962.

Schrider, D. R., and A. D. Kern, 2017 Soft sweeps are the dominant mode of adaptation in the human genome. Mol Biol Evol 34: 1863–1877.

Slatkin, M., 1985 Gene flow in natural populations. Ann Rev Ecol Systemat 16: 393–430.

Slatkin, M., 1991 Inbreeding coefficients and coalescence times. Genet Res 58: 167–175.

Stankowski, S., M. A. Chase, A. M. Fuiten, P. L. Ralph and M. A. Streisfeld, 2018 The tempo of linked selection: Rapid emergence of a heterogeneous genomic landscape during a radiation of monkeyflowers. BioRxiv.

Stankowski, S., J. M. Sobel and M. A. Streisfeld, 2017 Geographic cline analysis as a tool for studying genome-wide variation: a case study of pollinator-mediated divergence in a monkeyflower. Mol Ecol 26: 107–122.

Stankowski, S., and M. A. Streisfeld, 2015 Introgressive hybridization facilitates adaptive divergence in a recent radiation of monkeyflowers. Proc R Soc B-Biol Sci 282: 20151666.

Stephens, M., N. J. Smith and P. Donnelly, 2001 A new statistical method for haplotype reconstruction from population data. American Journal of Human Genetics 68: 978–989.

Stuart, Y. E., T. Veen, J. N. Weber, D. Hanson, M. Ravinet et al., 2017 Contrasting effects of environment and genetics generate a continuum of parallel evolution. Nat Ecol Evol 1: 0158.

Suchard, M. A., P. Lemey, G. Baele, D. L. Ayres, A. J. Drummond et al., 2018 Bayesian phylogenetic and phylodynamic data integration using BEAST 1.10. Virus Evol 4.

Tavares, H., A. Whibley, D. L. Field, D. Bradley, M. Couchman et al., 2018 Selection and gene flow shape genomic islands that control floral guides. Proc Natl Acad Sci U S A 115: 11006.

Team, R. C., 2016 R: A language and environment for statistical computing, pp. R Foundation for Statistical Computing, Vienna, Austria.

Terekhanova, N. V., M. D. Logacheva, A. A. Penin, T. V. Neretina, A. E. Barmintseva et al., 2014 Fast evolution from precast bricks: genomics of young freshwater populations of threespine stickleback Gasterosteus aculeatus. PLoS Genet 10: e1004696.

Troth, A., J. R. Puzey, R. S. Kim, J. H. Willis and J. K. Kelly, 2018 Selective trade-offs maintain alleles underpinning complex trait variation in plants. Science 361: 475–478.

Turner, T. L., M. W. Hahn and S. V. Nuzhdin, 2005 Genomic islands of speciation in Anopheles gambiae. PLoS Biol 3: 1572–1578.

Van Belleghem, S. M., C. Vangestel, K. De Wolf, Z. De Corte, M. Möst et al., 2018 Evolution at two time frames: Polymorphisms from an ancient singular divergence event fuel contemporary parallel evolution. PLoS Genet 14: e1007796.

Vijay, N., M. Weissensteiner, R. Burri, T. Kawakami, H. Ellegren et al., 2017 Genomewide patterns of variation in genetic diversity are shared among populations, species and higher-order taxa. Mol Ecol 26: 4284–4295.

Wallbank, R. W., S. W. Baxter, C. Pardo-Diaz, J. J. Hanly, S. H. Martin et al., 2016 Evolutionary novelty in a butterfly wing pattern through enhancer shuffling. PLoS Biol 14: e1002353.

Whitlock, M. C., and N. H. Barton, 1997 The effective size of a subdivided population. Genetics 146: 427–441.

Wright, S., 1931 Evolution in Mendelian populations. Genetics 16: 97–159.

Wu, T. D., and S. Nacu, 2010 Fast and SNP-tolerant detection of complex variants and splicing in short reads. Bioinformat 26: 873–881.

Yeaman, S., and M. C. Whitlock, 2011 The genetic architecture of adaptation under migration-selection balance. Evolution 65: 1897–1911.

